# Differential cell signaling testing for cell-cell communication inference from single-cell data by dominoSignal

**DOI:** 10.1101/2025.05.02.651747

**Authors:** Jacob T. Mitchell, Orian Stapleton, Kavita Krishnan, Sushma Nagaraj, Dmitrijs Lvovs, Christopher Cherry, Amanda Poissonnier, Wes Horton, Andrew Adey, Varun Rao, Amanda Huff, Jacquelyn W. Zimmerman, Luciane T. Kagohara, Neeha Zaidi, Lisa M. Coussens, Elizabeth M. Jaffee, Jeniffer H. Elisseeff, Elana J. Fertig

**Affiliations:** Department of Oncology, Sydney Kimmel Comprehensive Cancer Center, Johns Hopkins University, Baltimore, Maryland, USA; Johns Hopkins Convergence Institute, Johns Hopkins University, Baltimore, Maryland, USA; Johns Hopkins Bloomberg Kimmel Institute, Johns Hopkins University, Baltimore, Maryland, USA; Quantitative Sciences Division, Department of Oncology, Johns Hopkins University, Baltimore, Maryland, USA; Department of Genetic Medicine, Johns Hopkins University, Baltimore, Maryland, USA; Department of Biomedical Engineering, Johns Hopkins University, Baltimore, Maryland, USA; Translational Tissue Engineering Center, Johns Hopkins University, Baltimore, Maryland, USA; Institute for Genome Sciences, University of Maryland School of Medicine, Baltimore, Maryland, USA; Department of Medicine, University of Maryland School of Medicine, Baltimore, Maryland, USA; Department of Cell, Developmental & Cancer Biology, Oregon Health & Science University, Portland, Oregon, USA; Department of Molecular & Medical Genetics, Oregon Health & Science University, Portland, Oregon, USA; Knight Cancer Institute, Oregon Health & Science University, Portland, Oregon, USA; Department of Chemical and Biomedical Engineering, Johns Hopkins University, Baltimore, Maryland, USA; Department of Applied Mathematics & Statistics, Whiting School of Engineering, Johns Hopkins University, Baltimore, Maryland, USA; Greenbaum Comprehensive Cancer Center, University of Maryland School of Medicine, Baltimore, Maryland, USA; Institute for Health Computing, University of Maryland School of Medicine, Baltimore, Maryland, USA

## Abstract

Algorithms for ligand-receptor network inference have emerged as commonly used tools to estimate cell-cell communication from reference single-cell data. Many studies employ these algorithms to compare signaling between conditions and lack methods to statistically identify signals that are significantly different. We previously developed the cell communication inference algorithm Domino, which considers ligand and receptor gene expression in association with downstream transcription factor activity scoring. We developed the dominoSignal software to innovate upon Domino and extend its functionality to test statistically differential cellular signaling. This new functionality includes compilation of active signals as linkages from multiple subjects in a single-cell data set and testing condition-dependent signaling linkage. The software is applicable for analysis of single-cell data sets with multiple subjects as biological replicates as well as with bootstrapped replicates from data sets with few or pooled subjects. We use simulation studies to benchmark the number of subjects in compared groups and cells within an annotated cell type sufficient to accurately identify differential linkages. We demonstrate the application of the Differential Cell Signaling Test (DCST) in the dominoSignal software to investigate consequences of cancer cell phenotypes and immunotherapy on cell-cell communication in tumor microenvironments. These applications in cancer studies demonstrate the ability of differential cell signaling analysis to infer changes to cell communication networks from therapeutic or experimental perturbations, which is broadly applicable across biological systems.

## Introduction

Many algorithms for Cell-Cell Communication Inference (CCI)^1,2^ have been developed to infer how multicellular systems coordinate development and responses to stimuli via expression of genes encoding ligands and receptors from single-cell RNA-seq (scRNA-seq) datasets. Most CCI methods infer communication between cell types on the basis of co-expression of ligands by sender cells capable of activating receptors expressed by recipient cells. Methods that derive a signaling score as the product of ligand and receptor gene expression by interacting cell types showed poor accuracy in constructing signaling networks when applied to data sets with orthogonal quantification of signaling interactions^1^. One source of the inaccuracy of these estimates may rise from the failure to consider the expression of other genes encoding components of multi-subunit ligands or receptors. More advanced methods have sought to improve accuracy by accounting for the expression of signaling co-factors or measures of intracellular signaling cascades initiated following receptor activation such as target gene transcription^3–12^. The result of CCI is an inferred network of communication between cell types representing communication taking place at the time of sample collection.

One goal of many single-cell analyses is the inference of cellular changes between experimental contexts or treatment perturbations. In these cases, researchers may also seek to infer differences between cellular communication. However, most CCI algorithms are designed to infer communication in a single sample rather than specifically compare changes in communication between measured systems. As hundreds to thousands of ligand-receptor signals between pairs of cell types in a data set, CCI algorithms often rank ligand-receptor signals between cell types by a signaling score metric^4,6^ or provide a list of ligand-receptor pairs contributing to signaling between cell types^13^. However, these measures of signaling often do not provide a measure of the signal’s variance among samples to quantify the significance of observed differences between conditions. Qualitative comparisons of signaling may be drawn between aggregate measures of signaling between cell types between conditions as differences in scoring metrics or lists of ligand-receptor signals, but quantitative conclusions about signaling differences require statistical tests comparing the extended signaling cascade including ligands, receptors, and downstream response genes.

We developed Domino^13^ as a network-based CCI algorithm that relates cell-cell communication from ligand-receptor signaling to the receptor activation of downstream transcription factors (TFs) in intracellular signaling. Previously, we demonstrated that the binary score of ligand-receptor interactions prioritized by Domino could be used to quantifying differences in intercellular communication between cohorts of samples from different treatments tested in an immunotherapy clinical trial^16^. The sensitivity of this approach to sample size and cellular abundances was not assessed in these studies. Further extensions to compare conditions when limited sample numbers are pooled or to determine the changes to downstream transcription factor are also needed. To solve these problems, we develop the dominoSignal software as an R/Bioconductor package innovating upon Domino. This package has additional functionalities for comparing signaling networks between experimental conditions, inferring signals that utilize ligands and receptors that function as heteromeric complexes, and flexibly utilizing alternative ligand-receptor databases.

Approaches for differential comparison of inferred signaling have been implemented in other CCI methods that derive individual signaling scores for each investigated ligand-receptor interaction. MultiNicheNet^17^ derives a prioritization score for ligand-receptor signals that differentially occur between conditions that considers a product of magnitude of differential expression, signaling target gene expression, and proportion of cells expressing the ligand and receptor. CellChat^4^ generates a null distribution of their ligand-receptor score by calculating the score after randomly assigning cells to the groups being compared hundreds of times and assessing where the derived score falls on the null distribution to yield a z-score and p-value. As dominoSignal infers signaling as binary states of linkage between cell types via ligand and receptor pairs and linkage of receptors to transcription factors within a signaling cascade, we developed a statistical test for comparison of binarized signaling linkages called the differential cell signaling tests (DCST). The DCST utilizes Fisher’s Exact Test to statistically test the dependence of a signaling linkage differentially occurring between two groupings of samples represented in a scRNA-seq data set. We use the DCST to compare simulated signaling networks in order to assess the performance of identifying true differential signals, and we use DCST on real scRNA-seq data sets to demonstrate its functionality and the biological insights that can be gleaned from investigation of differential cell signaling using dominoSignal.

## Results

### The linkage summary structure organizes inferred intercellular and intracellular signaling interactions to facilitate testing of differential signals

In this study, we seek to extend these approaches to differential cell-cell communication, using the method Domino that also estimates intracellular signaling as the foundation. Briefly, Domino infers signaling by identifying TFs with differentially high activity in each cell type in a data set. The correlation between these TFs and receptor gene expression is assessed on a dataset-wide basis to establish TF-receptor linkages. An intracellular linkage between the receptor and TF is established if the TF is among those enriched in the cell type and a sufficient percentage of the cells in that cell type express the receptor. The presence of the intracellular linkage constitutes active receipt of signaling via the receptor in that cell type. Expression of the receptor’s ligands is then assessed by visualization of mean scaled expression of the ligands by cell types in the data set. dominoSignal provides discrete indications of signaling activity through the collation of linkages in which cell types participate. The DCST leverages this discrete quantification as the basis to compare inferred changes in cell-cell communication and is now implemented as a generalizable software for differential signaling analyses in scRNA-seq data sets measuring multiple experimental conditions for comparison (Figure 1).

**Figure 1:**
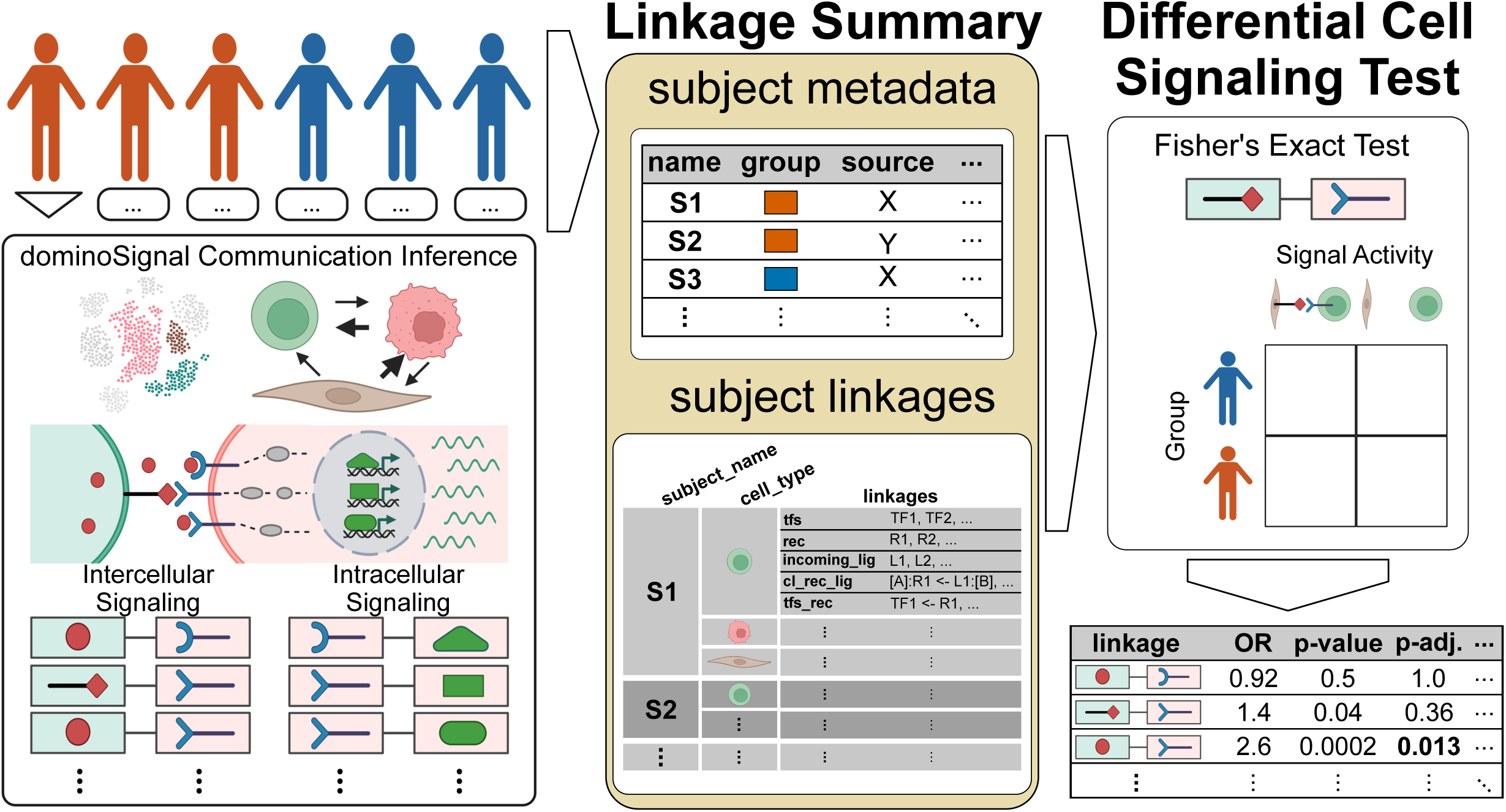
Framework of the differential cell signaling test (DCST). For a single-cell RNA-seq data set comprised of cells from multiple subjects, the cells are divided into groups based on the subject they originated from. Cells from each subject undergo cell-cell communication inference (CCI) with dominoSignal or other comparable CCI methods, resulting in lists of inferred intercellular ligand-receptor interactions grouped by recipient cell type as well as inferred intracellular signals between receptors and transcription factors or signaling targets if the CCI method permits. Inferred signals are organized in a data structure called a Linkage Summary consisting of a subject metadata table annotating the subjects considered and variables describing the subjects and subject linkages which is a nested list of subject names, the cell types present within the subject, and the inferred linkages from CCI which may include active features (tfs, rec, incoming_lig) and signaling features (intercellular: cl_rec_lig; intracellular: tfs_rec). The differential signaling test of a given linkage counts the occurrences of this linkage among the subjects included based on their assignment to a grouping variable to be compared across and tests for dependence of the linkage being present based on the grouping variable using a Fisher’s Exact Test. For each linkage, the odds ratio of the linkage occurring based on the grouping variable and the derived p-value are provided as an output as well as the p-value with false-discovery rate adjustment for multiple tests of linkages involving the same recipient cell type.

To implement differential cell signaling analysis, we developed a pipeline in which we first perform CCI for each sample in our scRNA-seq dataset independently. For example, if our dataset represents a cohort of tumors treated with different therapies, ligand-receptor network inference would first be performed for the data from each patient independently. This step ensures that inferred communication relies only on the multicellular environment for each sample. Signaling taking place between cell types via ligands and receptors is compiled in our software as **intercellular linkages**. In dominoSignal, these linkages are named following the format of:

[‘recipient cell type’]:‘receptor’ <- ‘ligand’:[‘sender cell type’]

Linkage naming initiates with the recipient cell because dominoSignal’s inference begins with evidence of receptor expression correlated with transcription factor activity in the recipient cell type. Network-based methods for CCI, including Domino^13^ and NicheNet^6^, further extend the network inference to estimate activation of TFs or individual target genes downstream of each receptor^18^. To account for inferred signaling between receptors and TFs or target genes, our software is also designed to compile these interactions as **intracellular linkages**. Changes in this cell-cell communication are then computed with a Fisher’s Exact Test designed to test the null hypothesis that the presence of an intercellular linkage between cell types is independent of a given comparative variable. In the example of multiple treatments described above, this analysis would employ Fisher’s Exact test on a contingency table of the number of subjects in each experimental group that include or lack the linkage being tested. Results are returned as a table including the odds ratio of the linkage being active in the reference group relative to the alternative group, the p-value derived from the Fisher’s Exact test, and FDR-adjusted p-values accounting for all interactions tested (Figure 1). This design based on the compilation of inferred linkages opens DCST to application with dominoSignal’s inferred networks or any CCI method that infers signaling on a cell type basis given there are criteria for a linkage to be on or off. Thus, a statistical basis for differential signaling across experimental conditions can be used to prioritize signals.

Interactions are encoded in dominoSignal by storing nested lists within a Linkage Summary object in a nesting hierarchy of subject identifier > receiver cell type > linkage type (Figure 1). Linkages are organized based on receiving cell type, following the recipient to sender direction of dominoSignal’s CCI approach. The linkage summary object also stores a table of meta data annotating features of each subject across which their signaling interactions can be compared to facilitate the implementation of the Fisher’s Exact Test for differential communication. The Linkage Summary more efficiently uses computer memory compared to maintaining Domino data objects for each of the samples as these objects also contain large matrices of gene expression data, calculated correlation coefficients, and ligand-receptor pairing references. The Linkage Summary’s simple formatting also allows for its use with other CCI methods that can be format inferred signaling as discrete linkages. This new infrastructure for differential cell signaling analysis is implemented in the current version of dominoSignal on Bioconductor.

### Signaling Simulation demonstrates sample sizes of at least 15 and cell types counts of at least 150 cells are sufficient for accurate identification of differential signals

We analyzed cell signaling changes using simulated data with a known ground truth to benchmark the performance of our differential signaling test under varying conditions. The simulated data were designed to benchmark performance of both identification of true positive signaling differences and avoiding detection of false positives. Briefly, we simulated single-cell data measuring two cell types (A or B). Cells within each cell type were assigned a probability of expressing two possible ligands (L1 and L2) and two possible receptors (R1 and R2). L1 and R1 are part of a signaling pair capable of linkage as are L2 and R2. We simulate simplified intercellular signaling with two simulation conditions (C1 and C2) where L1 – R1 signaling from A to B ([B]:R1 <- L1:[A]) differentially occurred in C1, L1 – R1 autocrine signaling from B to B ([B]:R1 <- L1:[B]) differentially occurred in C2, and L2 – R2 signaling from A to B ([B]:R2 <- L2:[A]) occurred in both conditions (Figure 2A).

**Figure 2:**
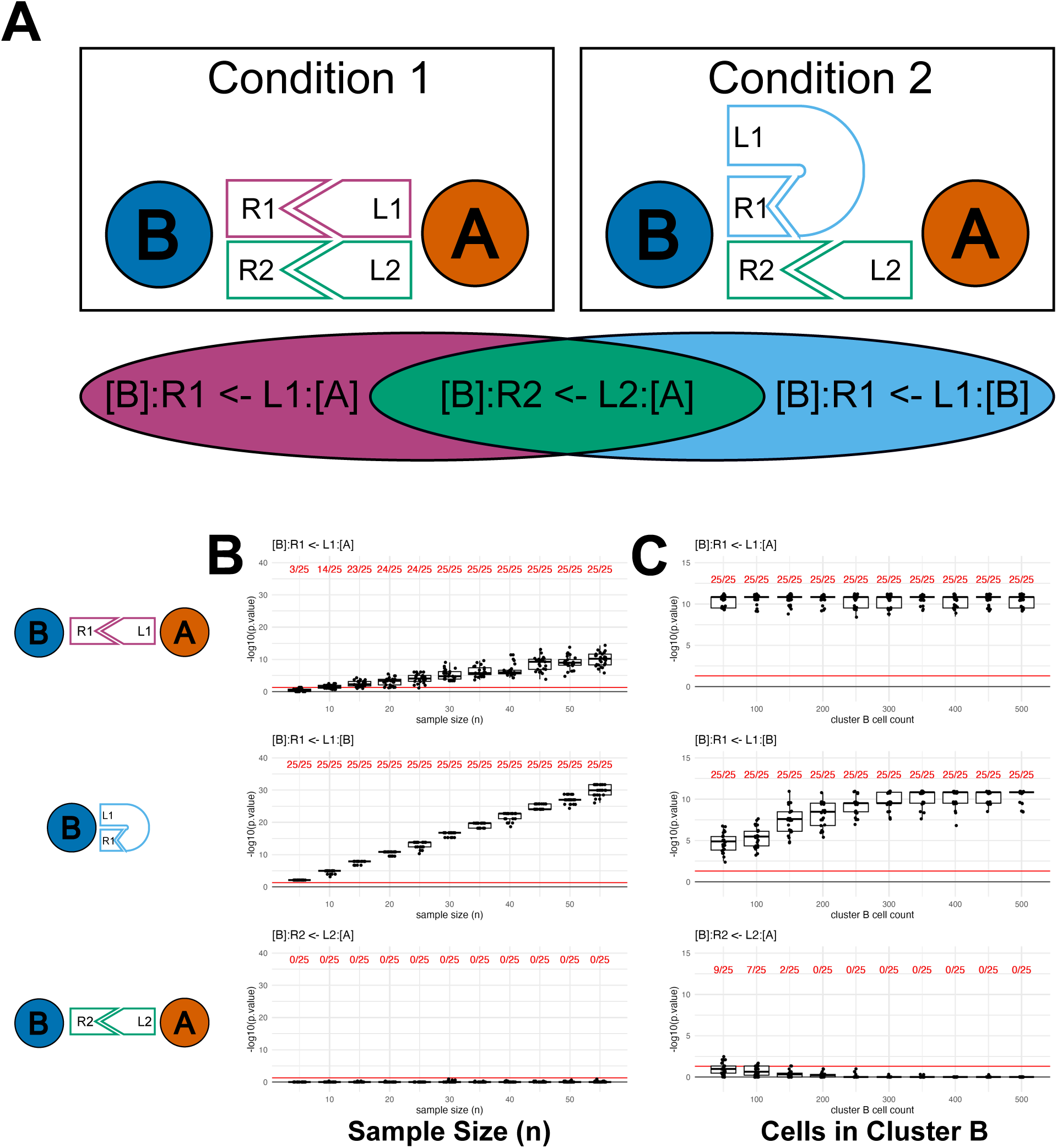
Assessment of performance of the differential cell signal testing in simulated data. (A) Graphical outline of signaling simulation. Cells belong to type A or B. Cells are simulated in two conditions with differing probabilities of signaling. Signaling occurs based on the percentage of cells expressing ligand-receptor pairs L1 to R1 and L2 to R2. The Venn diagram summarizes the interactions taking place exclusively in condition 1 (magenta), exclusively in condition 2 (blue), or equally likely in both conditions (green). Signals are phrased in terms of [“recipient cell”]:“receptor” <- “ligand”:[“sender cell”]. (B-C) P-values of differential signaling tests by Fisher’s Exact Test across repeated initializations of simulated data varying the number of subjects in conditions being compared (B) or number of cells belonging to type B (C). The tested intercellular signals are [B]:R1 <- L1:[A] (top), [B]:R1 <- L1:[B] (center), and [B]:R2 <- L2:[A] (bottom). The red line denotes a significance threshold of α = 0.05. The red proportion above denotes the number of unique initializations in which the tested achieved a p-value below 0.05.

Linkage criteria were simplified to ligand and receptor expression above a threshold to maintain control of expected results. A linkage was active if 25% of the cells in the sender cell type, either A or B, expressed the linkage’s ligand and 25% of the cells in the receiver cell type expressed the linkage’s receptor. A full table of starting parameters for the number of cells of each type in the two conditions is available as Supplemental Table S1. Both R1 and R2 had a 30% probability of expression by cells in cluster B in C1 and in C2. [B]:R1 <- L1:[A] was programmed to be a differential linkage more likely to occur in C1 by L1 having a 30% probability of expression by cluster A cells in C1 but a 25% probability of expression by cluster A cells in C2. [B]:R1 <- L1:[B], the autocrine signal, was programmed to be more likely to occur in C2. Cells in Cluster B had a 10% chance of expressing L1 in C1 and a 30% chance of expressing L1 in C2. [A]:R2 <- L2:[B] served as a negative control, as the linkage was equally likely to occur in C1 or C2. Cells in cluster A had a 30% chance of expressing L2 in both conditions. The first two linkages, [B]:R1 <- L1:[A] and [B]:R1 <- L1:[B], were assessed as true positives. Occurrences of [A]:R2 <- L2:[B] being found differential were false positives.

To determine the number of samples sufficient to identify true positive differential signals between conditions, simulations were conducted varying the number of subjects in each condition from 5 to 65 in increments of 5. For each parameter set, 25 unique initializations of the simulation were conducted, and the number of initializations where each of the linkages was significantly differential (p < 0.05) was counted (Figure 2B, Supplemental Table S2). A sample size of 30 was required to consistently identify [B]:R1 <- L1:[A] as differential across all 25 initializations, though a sample size of 15 sufficed to correctly identify the differential linkage 23/25 times. The other differential linkage, [B]:R1 <- L1:[B], was correctly identified across all sample sizes. The relative robustness of [B]:R1 <- L1:[B] being correctly identified as differential compared to [B]:R1 <- L1:[A] is likely due to the difference in L1 expression probability across conditions. The probability of L1 expression by cells in cluster B differed by 0.2 between conditions while the difference in L1 expression probability by cells in cluster A was 0.1. The negative control [B]:R2 <- L2:[A] linkage was correctly identified as having an equal probability of occurring in conditions C1 and C2 across all initializations. Based on these simulations, we estimate a minimum of 15 samples are needed to accurately infer differential communication, suggesting that these methods are accurately employed in cohort level atlas studies or require an alternative approach to overcome small sample sizes that are more common for preclinical studies.

Another challenge in analysis of scRNA-seq data is accuracy of data derived from rare cell types representing a small number of cells profiled in a scRNA-seq data set. Low cell counts have been shown to impact accuracy of identifying differentially expressed genes^19^, and could have similar impacts on inferred cell communication derived from gene expression in these cell populations. To assess the impact of low cell numbers on identification of differential signaling, the number of cells in cluster B in condition C1 was varied from 50 cells to 500 cells in increments of 50 while maintaining consistent probability of ligand and receptor expression and comparing 20 subjects each from conditions C1 and C2 by DCST (Supplemental Table S3). The accuracy of identifying true positive differential linkages [B]:R1 <- L1:[A] and [B]:R1 <- L1:[B] was maintained across all cluster B sizes. However, the median p-value derived from testing the autocrine [B]:R1 <- L1:[B] linkage increased with lower cell numbers (Figure 2C). The p-values were never high enough to fail to reject the null hypothesis that the proportion of simulated subjects with an active [B]:R1 <- L1:[B] linkage was equal between conditions C1 and C2. However, lower cell numbers raised p-values derived from the DCST (Figure 2C). The [B]:R2 <- L2:[A] linkage programed to not be differential between conditions was prone to false rejections of the null hypothesis of equal proportions of active linkage in each condition at cell numbers of 150 and lower. This experiment demonstrates that rare cell populations with few cells represented in a scRNA-seq data set are at increased risk of false positive detection of differential signaling. These simulations suggest that results concerning receipt of signaling by cell types with cell numbers below 200 may be unreliable and must be accompanied by orthogonal experiments to validate signaling dependence on the tested variables in real datasets.

### DCST analysis of a cohort of PDAC tumors demonstrates that classical PDAC exhibits increased fibroblast growth factor signaling compared to basal PDAC, validated in independent spatial transcriptomics data

Once benchmarked on simulated data, we next sought to demonstrate the performance of our DCST methodology in dominoSignal to determine how transcriptional phenotypes can alter intercellular signaling. Transcriptomic studies of pancreatic ductal adenocarcinoma (PDAC) by Moffitt et al^20^ described molecular subtypes of PDAC known as classical, characterized by increased cellular differentiation, and basal-like, characterized by greater plasticity and invasion potential. We expected that these differences in intrinsic properties of the neoplastic cells in a PDAC tumor would have ramifications on their interactions with non-neoplastic cells that constitute the PDAC microenvironment. To investigate this hypothesis, we obtained scRNA-seq data from the public data sets of Steele et al^21^ and Peng et al^22^, representing cells from 40 primary PDAC tumors. Cells were annotated according to Guinn et al^23^. We focus our analysis on epithelial cancer cells, CD8+ T cells, and fibroblast cell types (Figure 3A). Each tumor sample was classified into the subtypes defined by Moffitt et al^20^ based on the proportion of epithelial tumor cells of each subtype, resulting in 24 classical tumors and 16 basal tumors (Figure 3B, Supplemental Figure S1A-D). Our Differential CCI analysis was then applied based on these cellular and sample labels to demonstrate how this method could identify distinctions in basal and classical PDAC tumors in terms of how cancer cells interact with fibroblasts to shape the desmoplasia characteristic of PDAC and how CD8 T cells are hindered from cell killing by signaling withing the PDAC tumor microenvironment.

**Figure 3:**
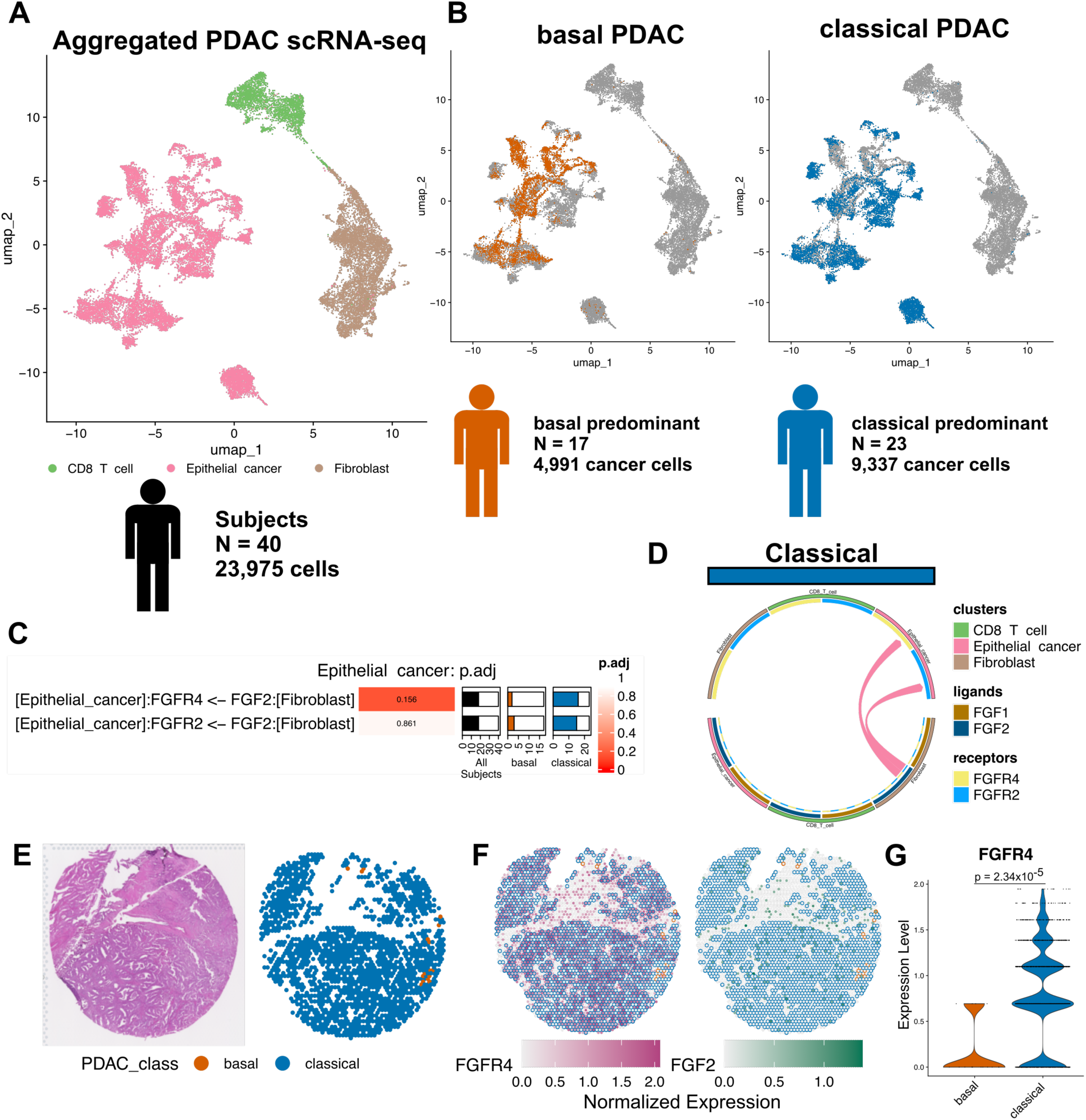
Consequences of DCST analysis comparing signaling between classical or basal PDAC subtypes. (A) UMAP of PDAC tumor cells scRNA-seq profiles from 40 subjects from Peng et al^22^ and Steele et al^21^ as annotated in Guinn et al^23^ as CD8 T cell (green), epithelial cancer (pink), and fibroblast (brown). (B) Typing of epithelial cancer cells as basal (orange) or classical (blue) based on gene module scores for gene sets described by Moffitt et al. Among subjects, 17 subjects had basal cells in the majority and 23 subjects have classical cells in the majority. (C) Leading differential signals from fibroblasts to epithelial cancer cells in classical or basal tumors. (D) Circos plot displaying differential signals via *FGF1* or *FGF2* to receptors *FGFR2* or *FGFR4* occurring in classical tumors. (E) Identification of classical (blue) and basal (orange) PDAC in Visium spatial transcriptomics data following the same module score approach employed on the scRNA-seq data. H&E image of PDAC tissue (left) is shown alongside Visium spots comprised of carcinoma cells colored by PDAC subtype (right). (F) Visualization of ligand and receptor expression in Visium spots where the spot center is colored by *FGFR4* (magenta, left) or *FGF2* (green, right) normalized expression and the border is colored based on the spot being classical PDAC (blue), basal PDAC (orange), or a non-carcinoma cell type (no border). (G) Violin plot of expression of *FGFR4* by basal PDAC (orange) and classical PDAC (blue) Visium spots. Overhead bar denotes the p-value from MAST test for differential gene expression.

Assessing differential signaling to epithelial cancer cells from fibroblasts identified 12 intercellular linkages differential between basal or classical subtypes before adjustment for multiple test correction (Supplemental Figure S1E, Supplemental Table S4). The top linkage inferred with this analysis was increased signaling of *FGF2* (Fibroblast Growth Factor 2) from fibroblasts to *FGFR4* (Fibroblast Growth Factor Receptor 4) on epithelial cancer cells in classical PDAC as compared to basal (FDR-adjusted p-value 0.156, Figure 3C-D). To validate that *FGF2* signaling is a distinguishing feature of classical PDAC from basal PDAC, we assessed co-expression of *FGFR4* and *FGF2* in a spatial transcriptomics data set of PDAC from Bell et al^25^ with co-occurrence of classical and basal PDAC cells within the same tumors. Among Visium (10X Genomics) spots annotated as cancer by Bell et al, spots were typed as classical or basal (Figure 3E). Cancer spots overlayed with *FGFR4* and *FGF2* gene expression showed a spatial absence of *FGFR4* expression in basal spots, demonstrating a diminished capacity to receive *FGF2* signals relative to classical PDAC (Figure 3F). Expression of *FGFR4* was significantly lower in basal cells than classical (Figure 3G, p-value = 2.34*10^-5^, MAST test^26^). Whether this *FGF2* signaling difference represents a point of therapeutic intervention to drive carcinoma cells towards or away from a basal de-differentiated state warrants further investigation.

### Bootstrapping enables robust detection of differential intercellular signaling in scRNA-seq data without biological replicates

The simulation studies and analysis of the human PDAC cohort demonstrate the use of DCST on scRNA-seq datasets containing multiple samples in each condition to be compared. Some studies limit sample sizes or pool biological replicates for cost-effective hypothesis-generation scRNA-seq studies. To extend the applicability of the DCST to small data sets lacking sufficient biological replicates to apply the Fisher test to compare between groups of samples, we developed an approach to simulating biological replicates through bootstrapping of cells. For each cell type in a data set, cells are sampled with replacement until the number of sampled cells matches the number of cells from that type in the parent sample, generating a bootstrap. Each generated bootstrap is then subjected to CCI, compiling the inferred intercellular and intracellular linkages within the Linkage Summary format to facilitate use of DCST (Figure 4A).

**Figure 4:**
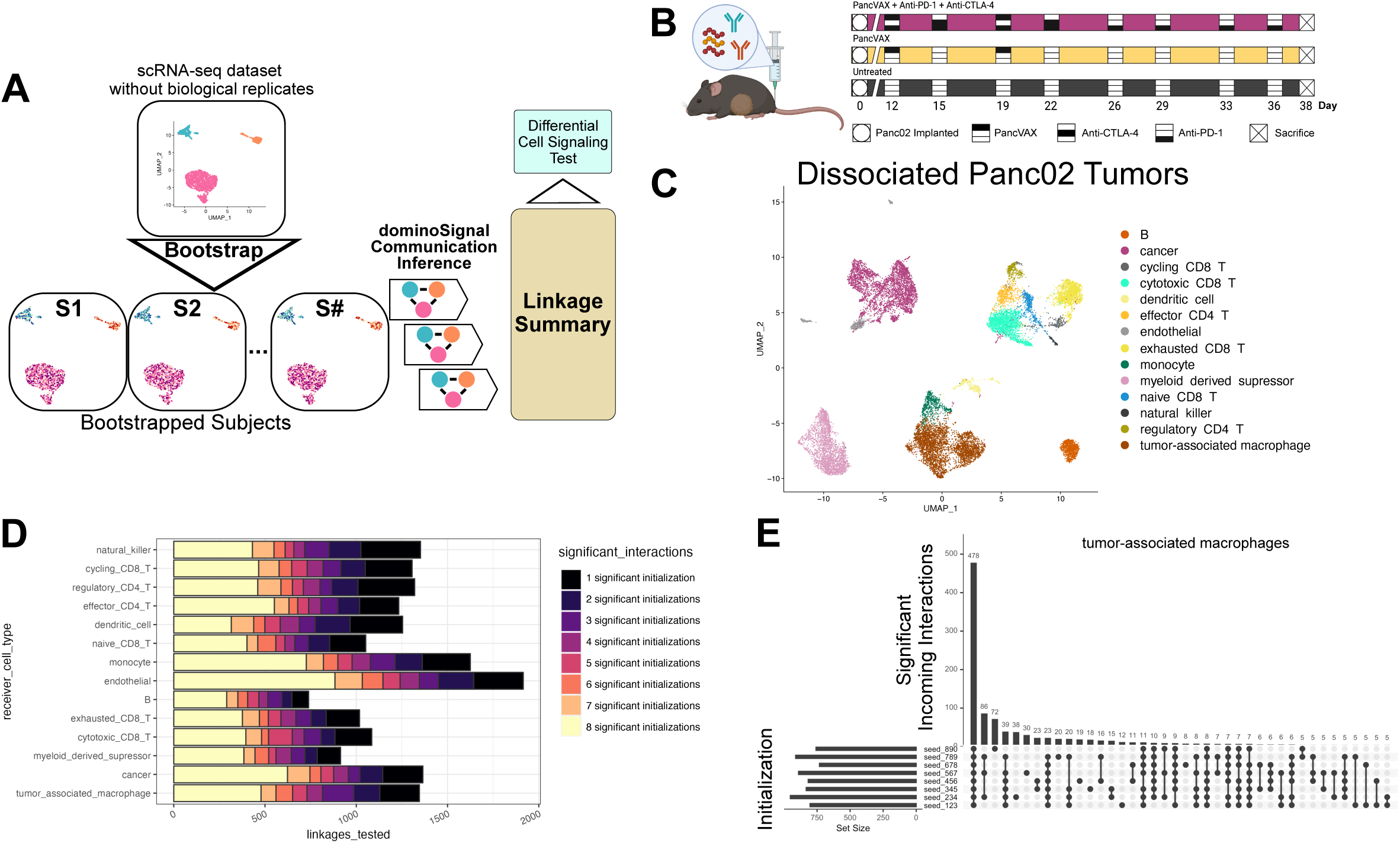
Assessing differential signaling in datasets without biological replicates through bootstrapping. (A) Graphical outline of bootstrapping approach in which cells are sampled with replacement from each cell type present in a subject until as many cells of that type from the original data set are obtained. Each bootstrapped sample is treated as an individual subject within the groups being compared via the differential signaling test. (B) Treatment regimen of mice which were implanted with Panc02 tumors and treated with PancVAX neoantigen cancer vaccine (gold), PancVAX in combination with anti-PD-1 and anti-CTLA-4 immune checkpoint inhibitors (magenta), or untreated given only isotype control (grey). (C) UMAP plot of scRNA-seq data of treated Panc02 tumors from Huff et al.^30^that is used as the basis of our bootstrapped DCST analysis. (D) Robustness of differential signals received by cell types between untreated and PancVAX-treated mice across eight unique initializations of bootstrapping. Bar plots display the number of tested linkages found to be differential in all eight initializations (yellow) down to those differential in only one initialization (black). (E) Upset plot counting the occurrence differential signals received by tumor-associated macrophages across the eight initializations.

We demonstrated this bootstrapping approach to DCST using a scRNA-seq data set designed to compare changes in the microenvironment between different therapeutic conditions. Briefly, we used scRNA-seq data from Huff et al^30^ that used scRNA-seq data to compare the tumor microenvironment changes from mice bearing subcutaneous tumors from the Panc02 cell line^31,32^ treated with either vehicle control (Untreated), a personalized neoantigen peptide vaccine (termed ‘PancVAX’), or PancVAX in combination with the immune checkpoint inhibitors anti-PD-1 and anti-CTLA-4 (PancVAX + anti-PD-1 + anti-CTLA-4) (Figure 4B). Profiling of these murine tumors revealed distinct expression profiles for multiple cell types critical to the tumor immune response including CD8+ T cells, CD4+ T cells, natural killer cells, B cells, macrophages, monocytes, myeloid suppressor cells, dendritic cells, endothelial cells, and cancer cells (Figure 4C).

We first sought to leverage simulations generated by distinct sampling of this dataset to benchmark performance of this bootstrapping approach prior to demonstrating the biological findings from these analyses. To ensure that bootstrapping consistency of inferred signaling, eight unique initializations of bootstrapping subjects each from the Untreated and PancVAX groups were conducted and subjected to DCST comparing the intercellular linkages received by each of the represented cell types. The number of initializations out of eight where each intercellular linkage was differential between Untreated and PancVAX was assessed (Figure 4D). For each cell type, at least half of the tested interactions were identified as differential, based on an FDR-adjusted p-value less than 0.05, in most of the initializations (>= 5/8 initializations). Among tested intercellular linkages received by tumor associated macrophages that were identified as differential in at least one initialization (Figure 4E), incoming linkages that occurred in all eight initializations were by far the most frequent (478/1346 tested linkages), though linkages identified as differential in single initializations were the second most frequent (217/1346 tested linkages). Using the same approach to assess robustness of received intercellular linkages in all cell types (Supplemental Figure S2), a majority of intercellular linkages identified as differential were robust across unique initializations of bootstrap sampling. However, cell types represented by very few cells, such as endothelial cells in this data set, have less consistency in the number of differential intercellular linkages across initializations. The occurrence of linkages found differential in a single initialization illustrates the risk of bootstrapping to overrepresent or underrepresent subpopulations of cells within a cell type that may be significant contributors to a CCI method’s criteria for intercellular linkage.

Variation generated through the bootstrapping approach could produce spurious identification of differential linkages caused by chance sampling of cells. We therefore tested if two unique initializations of bootstrapping from the same data set would result in any signals being identified as statistically differential by the DCST, despite being derived from the same parent data set. Differential signaling simulations had illustrated the importance of DCST comparisons including at least 15 subjects in each group being compared (Figure 2B). This established our basis that any bootstrapping from real data would use at least 15 bootstraps for each condition. Differential intercellular linkages were tested between two unique initializations bootstrapping from the PancVAX group (Supplemental Figure S3). No incoming intercellular linkages were found to be significantly differential between the two bootstrapping initializations from the same data. This demonstrates that variation generated through bootstrapping is not sufficient to cause false detection of differential linkages provided the number of bootstraps is large enough.

### Bootstrapped DCST analysis demonstrates that immune checkpoint inhibition halts immunosuppressive signaling from tumor-associated macrophages to exhausted CD8+ T cells in murine models of PDAC

Whereas the analyses above focused on leveraging our scRNA-seq data of immunotherapy combinations to benchmark algorithmic performance, previous studies demonstrated that the combination of PancVAX personalized vaccine with immune checkpoint inhibition is necessary to improve the survival of mice bearing Panc02 tumors^33^. Mechanistic studies further demonstrated that this improvement in survival is attributable to reinvigoration of cytotoxic effector functions in CD8 T cells expressing exhaustion markers^30^. Though the changes in gene expression by exhausted CD8 T cells between PancVAX and PancVAX + anti-PD-1 + anti-CTLA-4 have been assessed, intercellular signaling instigating the gene expression changes in exhausted CD8 T cells have yet to be identified. Thus, we sought to leverage our new DCST method and single-cell data to determine if the inclusion of anti-PD-1 and anti-CTLA-4 in a therapy regiment with PancVAX led to differential receipt of intercellular linkages by exhausted CD8 T cells as compared to PancVAX alone.

Bootstraps were generated from PancVAX and PancVAX + anti-PD-1 + anti-CTLA-4 cells and subjected to DCST of intercellular linkages for each recipient cell type (Figure 5A, Supplemental Table S5). The top 9 differential intercellular linkages from tumor-associated macrophages to exhausted CD8 T cells, more likely to occur in PancVAX over PancVAX + anti-PD-1 + anti-CTLA-4 (Figure 5B), revealed multiple immunosuppressive signals that diminished with addition of immune checkpoint inhibition. Among the differential signals sent by tumor-associated macrophages were interleukin-10 signaling (*Il10rb* <- *Il10*, *Il10ra* <- *Il10*), classically associated with suppressing cytotoxic T cell functions^34^, and apolipoprotein E signaling (*Sorl1* <- *Apoe*, *Ldlr* <- *Apoe*), demonstrated to be immunosuppressive in the context of pancreatic cancer^35^. *Cd80* <- *Cd274* signaling from macrophages, where *Cd274* encoding Programmed Death Ligand 1 (PD-L1), had greater occurrence in PancVAX over PancVAX + anti-PD-1 + anti-CTLA-4, an expected consequence of the anti-PD-1 therapy blocking PD-L1 to PD-1 signaling. Signaling to *Cd28* via ligands *Cd86* and *Icosl* was also more likely with PancVAX treatment. These signals are expected to be co-stimulatory to CD8 T cells^36,37^; however, this treatment had worse survival than PancVAX + anti-PD-1 + anti-CTLA-4^30^, indicating that the differential suppressive signals overwhelm the stimulatory signals. These results showcase the DCST’s applications in investigating signaling alterations caused by experimental therapy. Use of DCST revealed additional signaling targets for therapeutic modulation towards enhancing anti-tumor immunity. Such insights are made possible by extension of this test to account for pooled samples.

**Figure 5:**
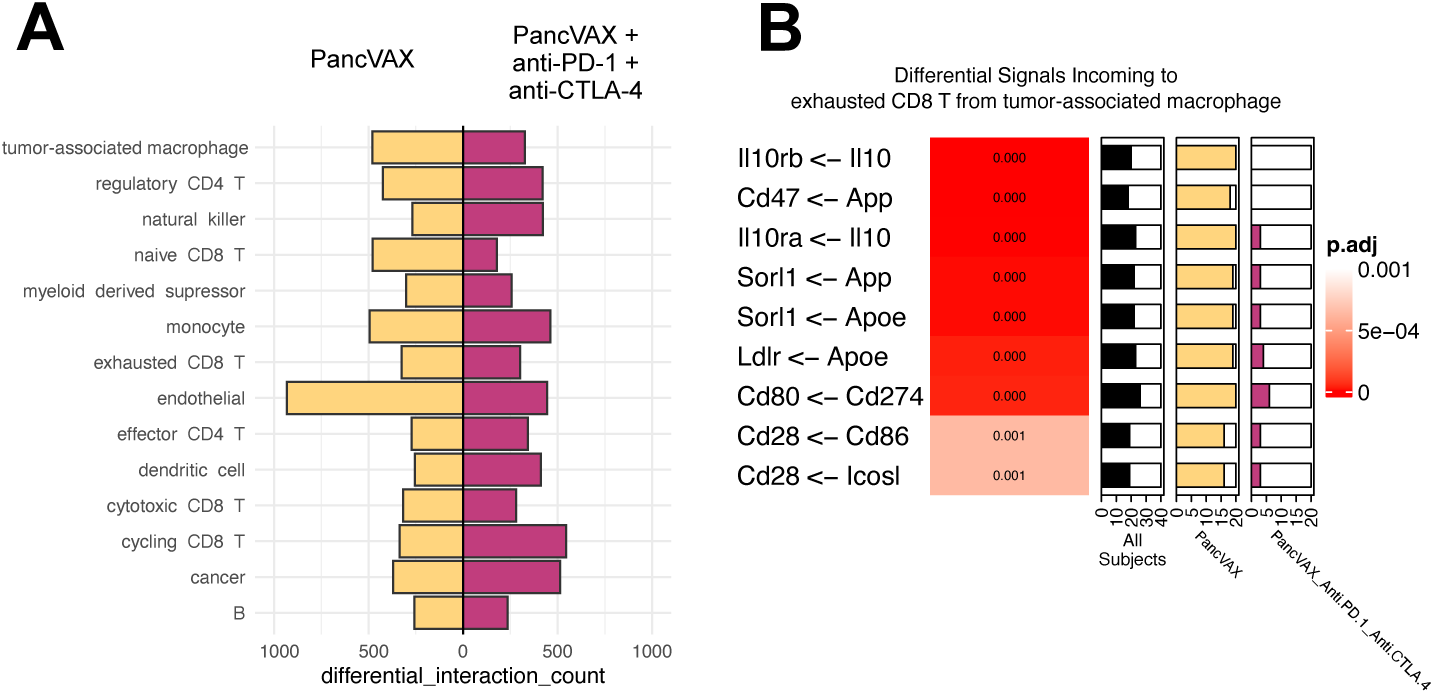
Immunosuppressive signaling by TAMs towards exhausted CD8 T cells is diminished in PancVAX + anti-PD-1 + anti-CTLA-4 treatment compared to PancVAX. (A) Counts of differential received signals in each cell type when comparing bootstrapped samples from PancVAX (n = 20, gold) to PancVAX + anti-PD-1 + anti-CTLA-4 (n=20, magenta). (B) Top nine differential signals from tumor-associated macrophages to exhausted CD8 T cells with higher probability of occurring in PancVAX over PancVAX + anti-PD-1 + anti-CTLA-4. Signals are phrased in terms of “‘receptor’ <- ‘ligand’”. Gradient values correspond to FDR-adjusted p-value. Proportion bars display the number of bootstrapped subjects with active signaling in all subjects (black), PancVAX (gold), or PancVAX + anti-PD-1 + anti-CTLA-4 (magenta).

### DCST analysis of integrated scRNA-seq and scATAC-seq data improves linkage testing specificity to demonstrate that histone deacetylase inhibition enhances macrophage inhibitory factor signaling in murine mammary carcinomas

The analyses to this point focused on demonstrating the performance of the DCST test for ligand-receptor pairs. An advantage of the dominoSignal CCI method is its further association of receptor activation to downstream estimates of TF activation using regulon scores computed with SCENIC^15,38^ and the intracellular linkage of these scores with receptor expression. SCENIC was selected for this estimation of TF activation as it refines the context-independent prior distribution of experimentally validated TF targets based upon the expression observed within an scRNA-seq data set. While expression is associated with TF activation, chromatin accessibility datasets provide more direct measure of the accessibility of DNA for binding to specifically infer context-specific TFs. Therefore, we sought to assess how our DCST tool infers downstream differences in TF activation and extend our pipeline to enable further analysis of single-cell Assay for Transposase Accessible Chromatin sequencing (scATAC-seq) to enhance those inferences. Recently, SCENIC was extended in a new tool called SCENIC+^39^ that enables direct refinement of regulons from scATAC-seq data and can be used directly as the source of TF activity in our dominoSignal software. As the integration of scATAC-seq had been shown by Gonzalez-Blas et al^40^ to enhance specificity of quantified TF activity, we sought to apply differential signaling testing to determine how a basis of TF activity with SCENIC or SCENIC+ affected inferred signaling by dominoSignal on experiments with paired scRNA-seq and scATAC-seq data.

To assess how the difference in TF activity quantification methodology would affect inferred signaling, we utilized scRNA-seq and scATAC-seq data reflecting mammary tumors collected from control MMTV-PyMT transgenic mice (UNT) and MMTV-PyMT mice treated with Entinostat (ENT), a selective class I histone deacetylase inhibitor, to rewire the immunosuppressive microenvironment of mammary tumors (Figure 6A)^41^. Cells were annotated as cancer associated fibroblasts (CAF), Lymphoid/Natural Killer (NK), Myeloid and Neoplastic based on marker gene expression or chromatin accessibility. Regulon activity was inferred from scRNA-seq data alone using SCENIC or paired scRNA-seq and scATAC-seq data using SCENIC+’s pseudo-multiome meta cell generation for each of the four cell types (Figure 6B, Supplemental Table S6). Among all 305 TF regulons quantified by the two methods in UNT cells, only 33 TFs were in common between the methods. In ENT, only 36 TFs were in common (Figure 6C). The much lower number of scored TFs agreed with previous results showing that SCENIC+ provides a more specific set of TF features than SCENIC by integrating chromatin accessibility^40^.

**Figure 6:**
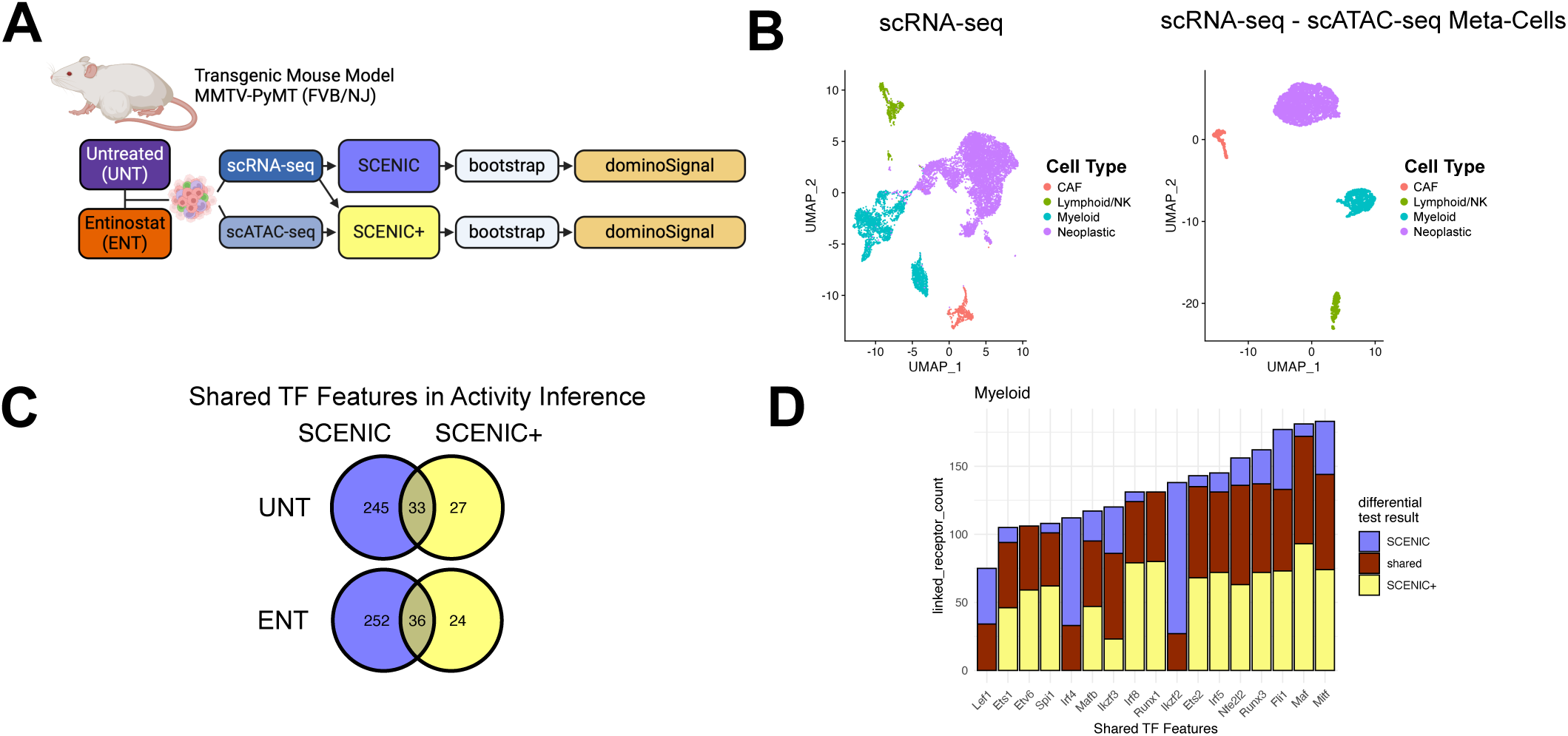
SCENIC+ multi-ome integration hones inference of intracellular linkages. (A) Experimental outline for multi-omics DCST analysis of data obtained from mammary carcinomas of MMTV-PyMT transgenic mice treated with Entinostat (ENT) or untreated controls (UNT). Dissociated tumors were subjected to scRNA-seq and scATAC-seq, and transcription factor activity was calculated with SCENIC on scRNA-seq data or with SCENIC+ on meta-cells using both scRNA-seq and scATAC-seq data. (B) UMAP representations of RNA-seq expression profiles (left) of dissociated tumors and SCENIC+-inferred meta-cell expression profiles (right). (C) Venn diagrams of quantified transcription factors from SCENIC (blue) and SCENIC+ (yellow) in the UNT and ENT treatment groups. (D) Stacked barplot counting receptors linked to transcription factors identified as active by dominoSignal in myeloid cells in the context of Entinostat treatment using SCENIC and SCENIC+. Intracellular linkages that were differentially identified when using SCENIC (blue), SCENIC+ (yellow), or equally likely between methods (shared, brown) were identified by DCST.

We then sought to determine if the inclusion of scATAC-seq data in our analysis impacted the intracellular linkages inferred by dominoSignal. Bootstraps were generated from the UNT scRNA-seq expression data and SCENIC TF activity scores to be compared to Bootstraps from the UNT meta-cell expression and SCENIC+ TF activity scores. For each cell type, dominoSignal identified TFs enriched in the cell type’s cluster and their linked receptors (Supplemental Figure S3). In myeloid cells, 17 TFs were found enriched using SCENIC or SCENIC+, and intracellular linkages involving these TFs were compared by DCST across methods. All TFs had intracellular linkages that were differentially inferred based on the usage of SCENIC or SCENIC+ (Figure 6D). For TFs that had differential linkages using both TF activity methods, more differential intracellular linkages were inferred while using SCENIC+. Though the SCENIC+ method narrows the number of TFs with quantifiable scores in a data set, there is not a decrease in the number of intracellular linkages from receptors to these TFs representing receptor regulation. dominoSignal’s inference of intracellular linkage is based on correlation between TF activity and receptor expression. Integration into meta-cells by SCENIC+ could favor higher measured correlations based on having fewer cells with expression drop out, thereby having more receptor-TF associations with sufficient correlation to be quantified as a linkage. SCENIC+’s demonstration of specificity favors the method’s use as the basis of CCI compared to SCENIC.

Based on the improved performance of the TF activity with scATAC-seq data, we proceeded to investigate the consequences of ENT treatment on communication between cell types in the MMTV-PyMT mammary tumor microenvironment using the SCENIC+ analysis. Upon assessment of differential intercellular linkages received by cell types with this integrated DCST analysis (Supplemental Tables S7-S9), receipt of macrophage migration inhibitory factor (MIF, encoded by *Mif*) by myeloid, lymphoid, and CAF cells were among the most likely to differentially occur in the ENT tumors (Figure 7A). MIF is a pleiotropic inflammatory cytokine that primes innate immune cells for activation^42^. MIF expression in MMTV-PyMT tumors has been associated with poor survival^43^. However, studies across multiple cancers indicate that the context of which cell type receives MIF signaling and what receptor interacts with MIF affect the functional outcomes of the signaling^44^.

**Figure 7:**
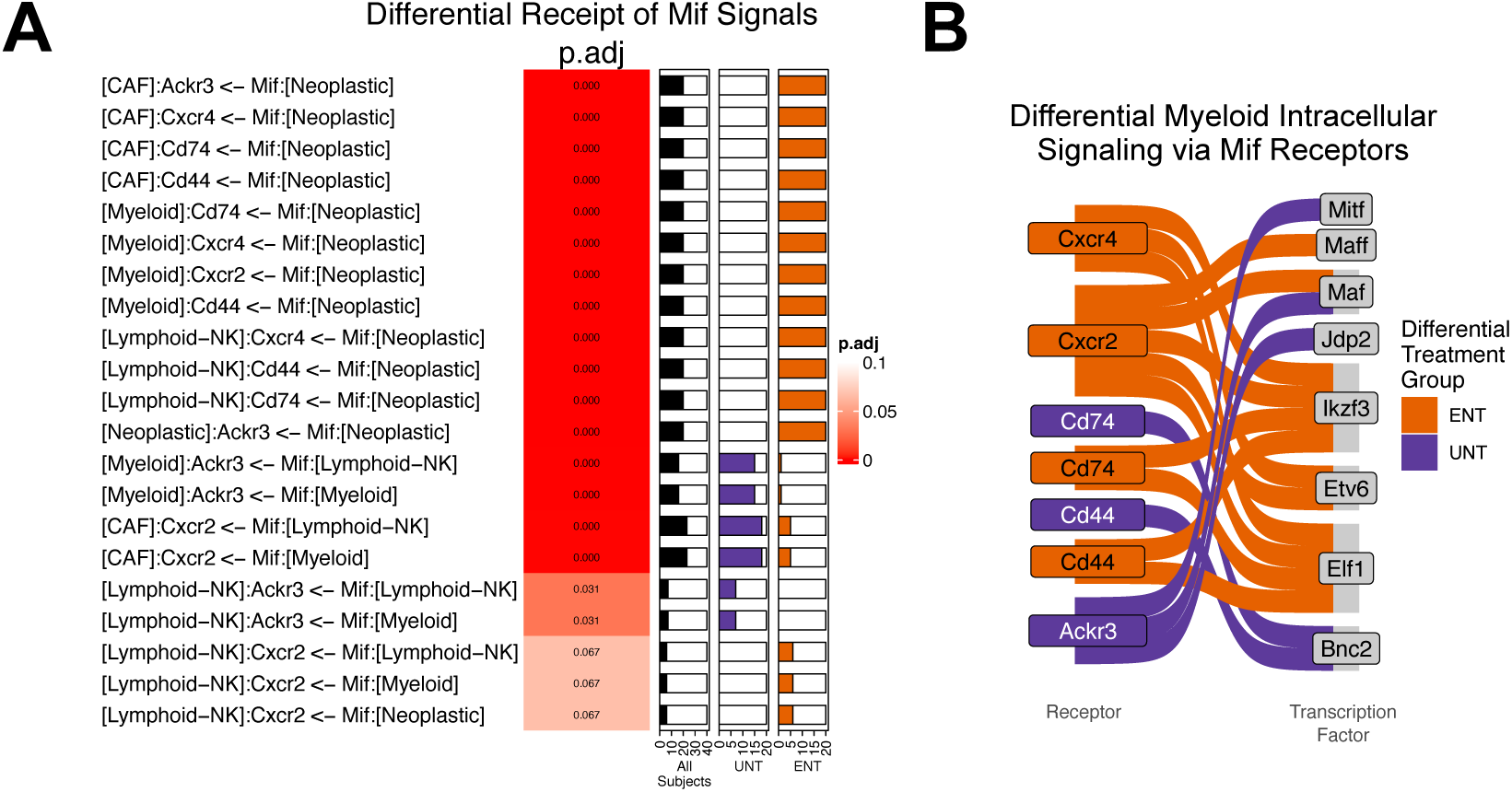
Entinostat enhances macrophage inhibitory factor signaling in MMTV-PyMT tumors. (A) Differential signaling via macrophage migration inhibitory factor (*Mif*) ligands between UNT- and ENT-treated tumors. (B) Sankey plots of differential intracellular signals between receptors of Mif ligands and transcription factors. Linkages are colored based on whether they are more likely to occur in UNT-treated (purple) or ENT-treated (orange) tumors.

Assessing all MIF signaling tested by DCST, the MIF intercellular linkages that differentially occur in ENT come from neoplastic cells towards CAFs, myeloid, and lymphoid cells via *Ackr3*, *Cxcr4*, *Cd74*, and *Cd44* receptors. Four intercellular linkages of lymphoid and myeloid cells signaling to myeloid cells via *Ackr3* and CAFs via *Cxcr2* differentially occurred in UNT (Figure 7A). CD74, encoded by *Cd74*, is the primary receptor of MIF, but signal transduction only takes place in the context of CD74 working in complex with the possible co-receptors CXCR2 (*Cxcr2*), CXCR4 (*Cxcr4*), CXCR7 (*Ackr3*), or CD44 (*Cd44*), which each having different outcomes of signaling^44^. To investigate the functional outcomes of these received signals, the intracellular linkages between MIF receptors and TFs was assessed by DCST in myeloid cells (Figure 7B). Among CD74’s chemokine co-receptors, CXCR2 and CXCR4 exhibited differential linkage in ENT tumors to TFs that drive lymphocyte development and function Ailos (*Ikzf3*) and E74 like ETS transcription factor 1 (*Elf1*)^45^, as well as Maf-F (*Maff*), which is activated as part of inflammatory signaling^46^. UNT tumors exhibited differential cell signaling via CXCR7 (*Ackr3*), which scavenges ligands away from CXCR4^47^, to *Mitf*, a TF that maintains cell survival^48^. These differential intracellular linkages to MIF receptors show that ENT histone deacetylase inhibitor treatment primes myeloid cells of the innate immune system for further activation by stimulating immune response transcription factors.

## Discussion

We present the DCST as a novel approach to testing differential signaling networks of inferred cell-cell communication in scRNA-seq data. We implement DCST using our CCI method Domino as a foundation and develop a new R/Bioconductor package dominoSignal for Domino and our DCST methods. dominoSignal includes a Linkage Summary data format for efficient storage of signaling inferred as intercellular and intracellular linkages on the basis of recipient cell type from many samples. The DCST provides a statistical basis for the comparison of these inferred signals within and between cell types across discrete independent variables in scRNA-seq data sets with many biological variables. We also include a bootstrapping approach for applications of DCST to data sets without annotation of biological variables to expand the contexts to which researchers can apply DCST.

We demonstrate through both simulated and real-world examples that the DCST method in dominoSignal distinguishes complex inter- and intra-cellular signaling changes between biological phenotypes and treatment conditions. Our simulated data examples required that signaling comparisons include at least 15 samples in each group being compared for consistent calling of true differential signals, with even larger sample sizes required to be powered for conditions with greater biological heterogeneity. We demonstrate that DCST results from bootstrapped samples are robust across unique initializations of bootstrapping, but identification of differential signals received by cell types represented by too few cells are more prone to inconsistent results. Our applications to real-world data show how the assessment of differential cell signaling can guide understanding of signaling interactions associated with distinct molecular states of pancreatic cancer, provide mechanistic insight into consequences of combination immunotherapy on networks of cellular interactions within tumors, and disentangle consequences of altered signaling via receptors with context-dependent signaling outcomes. We also use DCST to compare the methodologic basis of signaling networks inferred by dominoSignal when using SCENIC, RNA-seq-based, TF activity inference to dominoSignal using integrated scRNA-seq with scATAC-seq SCENIC+ TF inference.

Beyond DCST’s application to investigations of cancer biology and immunotherapy presented in this manuscript, comparisons of inferred signaling networks have broad applications in biological research. The DSCT method could have applications in identifying signaling alterations during organ development ^49^, comparisons of diseased and healthy tissues, and external perturbations to tissues. DCST leverages dominoSignal’s quantification of signaling in each sample as intercellular and intracellular linkages in binary on/off states to test dependence of these signals occurring upon a discrete sample variable. This binarization is unique amongst current approaches to comparing inferred signaling in scRNA-seq data^4,17^. The Linkage Summary format of storing incoming signals to cell types is agnostic of the method of inference. It could be employed to compare networks inferred by other CCI methods in addition to dominoSignal given that method-appropriate criteria of a signal being on or off are used.

We acknowledge that DCST is not appropriate for all comparisons of inferred signaling. DCST’s basis in Fisher’s Exact Test is well suited for comparisons of pairs of fixed states or groupings of samples. This test is not able to estimate effect sizes of incremental changes in continuous variables that describe samples. This could be important to investigations such as signaling consequences of increasing drug concentration or assumed additive effects of sample genotype. We also identified large sample sizes and cell counts within cell types as critical prerequisites to accurate identification of differential signals. Large sample sizes are often difficult to attain in scRNA-seq experiments due to financial cost^50^. Tests must also consider hundreds to thousands of incoming linkages to a cell type, necessitating multiple-test correction of the derived p-values. These considerations of sample size and cell number are best addressed by compendiums of multiple data sets, as demonstrated in this manuscript^23^, or large single-cell atlases representative of specific tissues or disease states^51^. For scRNA-seq experiments with small sample sizes or pooled samples, we provide a bootstrapping approach that uses observed data to estimate variation among biological replicates and demonstrated that this bootstrapping approach does not result in spurious identification of differential signals given that sufficiently large sample sizes are reached. This allows for the assessment of differential cell signaling in small-scale experiments. However, we acknowledge that bootstrapping results will better represent gene expression of the most ubiquitously expressed ligands and receptors while missing signals driven by rare cell populations within cell types.

DCST has the potential for wide application to compare signaling networks inferred in diverse experimental settings. Future directions of research may include the use of alternative statistical bases for comparing signaling interactions among samples. One such approach may be the use of logistic regression basis for DCST. The binary representation of an inferred signals activation state is well suited to dichotomous outcomes assessed by logistic regression. The generalized linear model basis of logistic regression allows for the consideration of covariate interaction effects on the signaling outcome^52^. However, even larger sample sizes are expected to be necessary to power usage of logistic regression compared to relatively small sample sizes of the non-parametric Fisher’s Exact Test. In addition to alternative statistical tests, criteria for the formatting of CCI results derived from other methods should be explored to enable applications of DCST to CCI methods beyond dominoSignal as well as direct comparisons of the signals identified by different CCI methods on the same data set.

In conclusion, we have developed the DCST as an extension to the dominoSignal R package for the statistical assessment of differential cell signaling between subjects in experimental groups. This package is available through Bioconductor with compatibility for standard formats of scRNA-seq analysis and applicable to multi-omics datasets including scATAC-seq data.

## Supplemental Figures

Supplemental Figure S1: Annotation of samples from PDAC compendium as basal or classical predominant. (A-C) UMAP plots of PDAC tumor cells scRNA-seq profiles from 40 subjects annotated by manuscript data source (A), basal module score (B), and classical module score (C). (D) Stacked barpot of samples colored by the percentage of epithelial cancer cells typed as basal (orange) or classical (blue). Bars are overlayed with percentage of cells belonging to the group. (E) Top 12 Differential intercellular signals between basal (orange) and classical (blue) subjects with raw p-values less than 0.05.

Supplemental Figure S2: Upset plots of the occurrence differential signals received cell types in Panc02 tumors across 8 initializations of comparing bootstraps from PancVAX and PancVAX + anti-PD-1 + anti-CTLA-4 treatment groups. Receiving cell types include B cells, cancer, cycling CD8 T cells, cytotoxic CD8 T cells, dendritic cells, effector CD4 T cells, endothelial cells, exhausted CD8 T cells, monocytes, myeloid-derived suppressor cells, naïve CD8 T cells, natural killer cells, regulatory CD4 T cells, and tumor-associated macrophages.

Supplemental Figure S3: Occurrence of differential intercellular linkages received by cell types from PancVAX data when comparing two initializations of generating 20 bootstraps from the same data.

Supplemental Figure S4: Stacked barplot counting receptors linked to transcription factors identified as active by dominoSignal in CAF, Lymphoid/NK, Myeloid, and Neoplastic cells in the context of Entinostat treatment using SCENIC and SCENIC+. Intracellular linkages that were differentially identified when using SCENIC (blue), SCENIC+ (yellow), or equally likely between methods (shared, brown) were identified by differential signaling test.

## Supplemental Tables

Supplemental Table S1: Simulation ligand expression probabilities

Supplemental Table S2: Simulation results when varying number of samples in each condition

Supplemental Table S3: Simulation results when varying number of cells in cell type B of condition 2

Supplemental Table S4: DCST results comparing intercellular linkages incoming to epithelial cancer cells between basal and classical samples

Supplemental Table S5: intercellular linkages incoming to exhausted CD8 T cells between PancVAX and PancVAX + anti-PD-1 + anti-CTLA-4 bootstraps

Supplemental Table S6: Counts of SCENIC+ meta-cell cell types

Supplemental Table S7: DCST results comparing intercellular linkages incoming to Myeloid cells between ENT and UNT bootstraps

Supplemental Table S8: DCST results comparing intercellular linkages incoming to Lymphoid-NK cells between ENT and UNT bootstraps

Supplemental Table S9: DCST results comparing intercellular linkages incoming to CAF cells between ENT and UNT bootstraps

## Methods

### DominoSignal for inference of inter- and intra-cellular signaling

dominoSignal is an R software package developed as an iteration upon the Domino R package developed by Cherry et al^13^. Domino conducts CCI in scRNA-seq data sets on the basis of inferring evidence of signal receipt based on Spearman correlation of genes encoding receptors with and a measure of TF activity in each cell. Ligand expression is then assessed based on the mean of scaled expression of the ligand-encoding genes by each cell type in the data set.

dominoSignal CCI on a scRNA-seq data set requires the data set’s raw RNA count matrix, the normalized and scaled RNA matrix, a TF by cell TF activity score matrix, and a table describing the possible ligand-receptor pairs called an rl_map. Optionally, researchers may include a character vector of cell assignments to clusters or cell types so that signal-receipt on a per-cell type basis may be assessed and a list of genes inferred to be regulatory targets of each TF as a “regulon” which can be used in the dominoSignal analysis to remove TF-receptor linkages that may be attributable to TF activation driving receptor expression. The basis of TF activity quantification is up to the discretion of the user. Conventionally, Domino and dominoSignal have used SCENIC^15^ as the basis of TF-activity quantification, but any method that results in a TF by gene matrix of activity scores is a valid input. The “rl_map” must include columns annotating the interacting ligand-receptor pairs. These include: int_pair, the names of the interacting ligand and receptor separated by “ & ”; gene_A, the gene or genes encoding partner A; gene_B, the gene or genes encoding partner B; type_A, (“L”, “R”) - indicates whether partner A is a ligand (“L”) or receptor (“R”); and type_B, (“L”, “R”) - indicates whether partner B is a ligand (“L”) or receptor (“R”). For ligands or receptors that function as heteromeric complexes encoded by multiple genes, the names of all genes in the complex are included in the gene_A or gene_B columns separated by commas. dominoSignal includes a helper function, *create_rl_map_cellphonedb*, to format database tables from CellPhoneDB^3^ annotating human ligand-receptor pairs into an rl_map format. rl_maps for mouse ligand-receptor interactions were generated from CellTalkDB^14^.

The first stage of the dominoSignal analysis is carried out by the *create_domino* function. This function takes the required data described above as inputs with additional parameters “use_clusters” which dictates if cell type assignments will be considered for receptor activation and ligand expression as well as “use_complexes” which dictates if heteromeric complex ligands and receptors included in the rl_map will be considered during CCI. The result of *create_*domino is a domino object, an S4 class that stores the input data used for CCI as well as information on the tests for signaling inference and the resulting signaling network. *create_domino* begins CCI by testing for enrichment of TF activity in each cell type by conducting a one-sided Wilcoxon rank-sum test of whether each TF’s mean score in the cell type is significantly greater than the mean score in all other cells. A matrix of p-values from the Wilcoxon rank-sum tests are stored in the “de” slot of the resulting domino object. Spearman correlations between the TF activities and receptor gene expression are then assessed across all cells in the data set. Calculated Spearman correlation coefficients are stored in the “cor” slot of the domino object. For complex receptors encoded by multiple genes, the stored correlation values are the median coefficients among all component genes. If a list of TF regulons is provided to the “tf_targets” argument to *create_domino*, receptor that is part of a TF’s regulon will be set to 0 correlation to avoid assigning linkage between a TF that drives a receptor’s expression rather than the receptor driving TF activity. With TF enrichment and receptor-TF correlations calculated and stored in the domino object, the next step pertains to setting parameters for assigning intracellular and intercellular linkages.

The next step of dominoSignal analysis is carried out by the *build_domino* function. This function takes the domino object resulting from create_domino as an argument as well as parameters for assigning linkages. “min_tf_pval” sets the maximum Wilcoxon rank-sum test p-value for assigning a TF as enriched in each cell type. A cell type’s enriched TFs are stored as a list within the domino object’s “linkages$clust_tf” slot. “rec_tf_cor_threshold” sets the minimum Spearman correlation coefficient between a TF and receptor to assign an intracellular linkage between them. Data set-wide TF-rec linkages are stored in the “linkages$tf_rec” slot. Intracellular linkages within cell types are assigned on the basis of enrichment of the TF in the cell type as well as non-zero expression of the receptor in a minimum percentage of cells within the cell type specified by the “min_rec_percentage” argument to *build_domino*. Active receptors for each cell type are stored in the “linkages$clust_rec” slot and their intracellular linkages with TFs is stored within “linkages$clust_tf_rec”. Intercellular linkages annotating a sender cell type, ligand, receptor, and receiving cell type can be annotated in “linkages$rec_lig_cl” based on mean scaled ligands expression above a threshold using the *add_intercellular_linkages* function where the threshold is specified by the “signal_threshold” parameters.

At this point, the domino object can be used for creating signaling plots presented in the original version of Domino as published by Cherry et al^13^ including TF feature expression by clusters, and heatmaps or networks of cumulative signaling between all clusters or specific linkages that each cluster takes part in. dominoSignal adds capabilities to render circos plots of the expression of ligands signaling to a receptor inferred to be active where the width of chords leaving cell type arcs corresponds to magnitude of ligand gene expression. Vignettes tutorizing the usage of dominoSignal, plotting dominoSignal results, navigating domino objects, and using SCENIC for TF activity quantification is available from https://fertiglab.github.io/dominoSignal/index.html using a reference scRNA-seq data set of 4000 peripheral blood mononuclear cells collected by 10X Genomics^53^.

### Parameters for dominoSignal cell-cell communication inference on real data sets

For all the real single-cell datasets, CCI was conducted with dominoSignal (v1.0.6). All TF activity scoring was conducted using SCENIC^15,38^ (v 0.11.0) or SCENIC+^40^ (v 1.0). Unless otherwise specified, all instances of using dominoSignal were conducted using parameters min_tf_pval = 0.001, rec_tf_cor_threshold = 0.15, min_rec_percentage = 0.05, and no limits on the number of TFs inferred as active in a cell type or receptors linked to a transcription factor. Cell types were identified as ligand senders in intracellular linkages if the mean scaled expression of the ligand was greater than 0.

### The Linkage Summary structure

The Linkage Summary is an S4 class object in the R computing environment defined within the dominoSignal R package to organize inferred cell-cell communication via ligands to receptors and intracellular associations of receptors with gene or TF targets of activation. The Linkage Summary object stores the linkages slot from multiple domino objects to facilitate their comparison and save space in computer memory by not including the other data in the domino objects such as expression matrices. The linkage summary consists of three slots. “subject_linkages” is a nested list organized on the hierarchy of subject > receiving cell type > linkages. Each subject’s slot in the list is populated with all linkages inferred by running dominoSignal on the cells from that subject. The character vector of intercellular linkages from the “rec_lig_cl” slot provide a binary representation of whether each possible interaction is on or off. The linkage summary also retains character vectors of intracellular linkages from the “clust_tf_rec” and all other signaling features stored in the linkages slot of each domino object. Though this is tailored to retrieving this information from a domino object, the Linkage Summary is agnostic to the method used to infer intercellular linkages. Thus, a Linkage Summary could be used to store and compare any inferred communication on the basis of cell type so long as the linkages of communication could be binarized as active or inactive. The Linkage Summary’s “subject_meta” slot contains a table listing the subjects with columns annotating variables describing the subjects that could be used for DCST comparisons. The Linkage Summary’s “subject_names” slot contains a factor vector of the names of subjects for easy access.

### Differential cell signaling test (DCST) functions

The dominoSignal package also includes functions for tabulating the active linkages among subjects covered in the linkage summary (*count_linkages*) and conducting a DCST for a specified linkage type received by a shared cell type across independent variables annotated in the subject_meta table (*test_differential_linkages*). The *test_differential_linkages* function populates a contingency table for the linkage being tested where rows are the levels of the independent variable being tested and columns are the linkage being inferred as active or inactive. The resulting table, where each row corresponds to a tested linkage, provides the name of the linkage feature, the total number of subjects (total_n) and number of subjects with the active linkage (total_count), and columns for each level of the independent variable counting the number of subjects in the level (‘level’_n) and number of subjects with the active linkage (‘level’_count).The DCST uses Fisher’s Exact Test for statistics and returns the odds ratio of signaling taking place among subjects in the reference group relative to signaling taking place in subjects of the alternative group and the derived p-value are provided. Adjusted p-values (p.adj) are calculated using Benjamini-Hochberg false-discovery rate correction as implemented in the p.adjust function in the R stats package (v 4.2.0). This adjustment accounts for all unique linkages incoming to the cell type across all subjects

### Simulation of cell-cell communication and communication inference

Simulated data were generated with predetermined intercellular linkages specified to be differential or consistent between conditions to have a ground truth against with to assess methods performance. The purpose of these simulations were to generate samples with expected intercellular linkages between two cell types. Ligand and receptor expression by cells was simplified to binary states of expressed (1) or not expressed (0) in each cell. The criterion for an intercellular linkage via these a ligand-receptor pairs was that at least 25% of the cells in the sender cell type express the ligand and that at least 25% of recipient cells express the receptor.

Two experimental conditions, C1 and C2, were designed for simulating cells for assessing intercellular linkages. Cells belonged to one of two cell types, A or B. Expression of two ligands, L1 and L2, and two receptors, R1 and R2, were simulated for each cell. L1-R1 and L2-R2 form pairs capable of intercellular linkage. When each cell was generated, the cell had a programmed probability of expressing each of the four signaling molecules based on a draw from a uniform distribution. These probabilities were specified based on the condition being simulated and the cell’s type. For example, a cell generated from cell type A in condition 1 had a 30% probability of expressing L1, 30% probability of expressing L2, 5% probability of expressing R1, and 5% probability of expressing R2. Probabilities of expression for each simulated cell type and condition under default parameters are listed in Supplemental Table S1. Upon generating all cells for a simulated sample, presence of intercellular linkages was assessed based on the intercellular linkage criteria stated above, and the result was stored as a Linkage Summary. Differential cell signaling between C1 and C2 was assessed by DCST using this Linkage Summary. The default parameters were designed so that the [B]:R1 <- L1:[A] linkage was more likely to occur in C1, [B]:R1 <- L1:[B] linkage was more likely to occur in C2, and [B]:R2 <- L2:[A] was equally likely to occur in each condition.

The effect of the number of subjects on correct inference of differential intercellular signaling was assessed by varying the number of subjects on identifying differential intercellular linkages for each condition from 5 subjects up to 65 subjects in increments of 5. The effect of cell number on correct inference of differential intercellular signaling was assessed by varying the number of cells in cell type B in condition C2 from 50 cells to 500 cells in increments of 50. The raw p-values derived from the DCST were compiled from 25 initializations of each parameter set and assessed for statistical significance at α = 0.05. Raw p-values were used as only 3 possible linkages were tested for each initialization.

### Bootstrapping of sample replicates for DCST analysis of pooled scRNA-seq data

In the cases of assessing DCST in scRNA-seq data sets without annotation of source subjects as biological replicates, variation representative of biological replicates was generated by bootstrapping. To bootstrap, the scRNA-seq data set was separated into the two groups to be compared by DSCT. For each bootstrap from a group, cells were generated on a per cell type basis by uniform sampling with replacement, as implemented by the sample function in the base package in R (v 4.2.0). Sampling took place until as many cells of the type were drawn as were present in the original group. For CCI by dominoSignal in bootstrapped data, cells in the bootstrap had the same TF activity scores, raw RNA counts, and normalized RNA expression as the corresponding cells from which they were sampled. RNA expression scaling was conducted on the bootstrapped normalized RNA expression once all sample draws were complete. CCI with dominoSignal on each bootstrap proceeded using parameters as described in ‘Parameters for dominoSignal cell-cell communication inference on real data sets’. Linkages inferred by dominoSignal were stored as a Linkage Summary where each bootstrap was treated as a unique subject and annotated with the condition of the original data used to generate the bootstrap in the “subject_meta” table.

### Single-cell RNA-seq data from human Pancreatic Ductal Adenocarcinoma tumors

Single-cell RNA-seq data from human PDAC tumors were obtained from the datasets compiled and annotated as described in Guinn et al^23^. In brief, this previous study integrated six data sets of scRNA-seq profiles from primary PDAC tumors^21,22,54–57^ and performed clustering to identify 11 cell types present in PDAC tumors^23^. In this current study, cells belonging to the fibroblast, epithelial cancer, and CD8 T cell types the two largest data sets used by Guinn et al^23^, 14,537 cells from Peng et al and 9,428 cells from Steele et al (Supplemental Figure S1A). These data were isolated into a Seurat object^58^ and batch-corrected PCA embeddings for cells correcting for manuscript source were calculated using Harmony^59^ prior to UMAP visualization^60^. Seurat module scores for gene sets associated with classical PDAC and basal-like PDAC as defined by Moffitt et al^20^ were calculated for all epithelial cancer cells (Supplemental Figure S1B). Cells were typed as classical or basal based on which of the two module scores were greater in the cell (Figure 3B). The 40 subjects from which cells were derived were annotated as basal or classical if 50% or greater of their epithelial cancer cells were of that cancer cell type (Supplemental Figure S1C). Intercellular linkages received by epithelial cancer cells were compared between basal and classical subjects using DCST. Intercellular linkages with p.adj values less than 1 (Figure 3C) and un-adjusted p-values less than 0.05 (Supplemental Figure S1D) were plotted as differential signaling plots from dominoSignal (v 0.2.1). Circos plots showing FGF-FGFR signals approaching statistical significance for differential linkage were generated using circlize (v 0.4.15).

### Analysis of Visium spatial transcriptomics of pancreatic ductal adenocarcinoma

Visium spatial transcriptomics data was obtained from a cohort of formalin-fixed and paraffin-embedded human surgical specimens containing pancreatic intraductal neoplasia proximal to PDAC tumors underwent spatial transcriptomics profiling using Visium spot-based capture (10X Genomics) as described in Bell et al^25^ and Lyman et al (In review). RNA counts matrix outputs from SpaceRanger were downloaded from NIH GEO repositories GSE254829 and GSE294669. In total, 20 tissue sections collected from five patients (GSE254829: PanIN low grade1 R, PanIN low grade2 R, PanIN low grade3 R, PanIN low grade4 R, PanIN low grade5 R, PanIN low grade6 R, PanIN high grade1 R; GSE294669: CP01, NRL01, PANIN01, PDAC01, NRL02, PDAC02, CP03, NRL03, PDAC03, PDAC04, CP05A, NRL05, PDAC05) underwent spot annotation of tissue type using the CODA machine learning tool for segmentation and annotation of cell types in hematoxylin and eosin images^25,61^ and quantification of PDAC basal-like subtype and classical subtype module scores^20,25^. Tissue section PDAC03 was selected for spatial validation of differential signaling identified in the human PDAC scRNA-seq data as the section had many spots containing PDAC (1696 PDAC spots). Spots typed as “PDAC” by CODA were further annotated as “basal” or “classical” based on which of the two scores in the spot were higher (Figure 3E). RNA Expression of *FGFR4* and *FGF2* normalized using the Seurat SCTransform function (v 4.1.1) was assessed for overlap with basal and classical regions of PDAC tissue and differential expression using the MAST test^26^ as implemented in Seurat^58^.

### Differential Cell Communication Testing on Panc02 Tumor Response to PancVAX and Immune Checkpoint Inhibitor Therapy

Single-cell RNA-seq data of vaccine and immune checkpoint inhibitor treatment of a subcutaneous, murine model of PDAC generated from the Panc02 cell line are obtained from Huff et al^30^ (GEO GSE244992). This dataset contains pooled cells from 5 mice from untreated controls, mice treated with the PancVAX^33^ personalized vaccine that consists of 12 major histocompatibility complex I-restricted peptides bearing mutations in the Panc02 cell line, anti-PD-1, anti-CTLA-4, and their combinations. Briefly, cells were annotated by clustering into B cells, cancer cells, cycling CD8 T cells, cytotoxic CD8 T cells, dendritic cells, endothelial cells, exhausted CD8 T cells, monocytes, myeloid-derived suppressor cells, naive CD8 T cells, natural killer cells, regulatory CD4 T cells, and tumor-associated macrophages according to the marker genes described in detail in Huff et al^30^.

In this current study, cells were subset to those belonging to the Untreated, PancVAX, and PancVAX + anti-PD-1 + anti-CTLA-4 treatment groups. Normalized expression was rescaled for cells in each treatment group using the Seurat ScaleData function. TF activity score inference was conducted on each treatment group independently using SCENIC^15,38^ (v 0.11.0) with mouse TF lists and TF motif references for the mm10 reference genome provided by cisTarget^62,63^ (https://resources.aertslab.org/cistarget/databases/). For comparison of bootstrapped DCST results between Untreated and PancVAX, 20 bootstraps were generated for each treatment group with eight unique seeds (123, 234, 345, 456, 567, 678, 789, 890). For comparison of signaling by DCST between PancVAX and PancVAX + anti-PD-1 + anti-CTLA-4, 20 bootstraps were generated for each treatment group with the seed 123. CCI was conducted using dominoSignal (v 0.2.1) on each bootstrap. Parameters for dominoSignal to infer linkages were consistent with those described in the ‘Cell-Cell Communication Inference by dominoSignal’ section of the methods. Ligand receptor pairs were sourced from CellTalkDB^14^ (v 1.0), and possible interaction between ligand-receptor pairs *Cd80*-*Ctla4* and *Cd86*-*Ctla4* were manually added to the reference as they were not included in CellTalkDB v1.0 and are expected to be a target of anti-CTLA-4 therapy.

Intercellular linkages were considered significantly differential across treatments if the p-value derived from the Fisher’s Exact Test after FDR correction accounting for intercellular linkages incoming to the recipient cell type tested was less than 0.05. Benjamini-Hochberg FDR adjustment was conducted with the p.adjust function in the R stats package (v 4.2.0). Stacked barplots of the occurrence of differential intercellular linkages was made with ggplot2 (v 3.4.1). Differential signaling plots were made with dominoSignal (v 0.2.1).

### scRNA and scATAC-seq profiling of MMTV-PyMT murine mammary tumors treated with Entinostat

MMTV-PyMT mice, which develop mammary adenocarcinoma tumors due to expression of the *PyMT* transgene under an MMTV promoter, were used as a representative sample of mammary cancer with minimal cytotoxic immune cell infiltrates. At 80 days of age, female mice were randomized into assignment for treatment with Entinostat (ENT), an oral histone deacetylase inhibitor, delivered in mouse chow for a targeted oral intake of 12 mg/kg or assignment for untreated control (UNT) receiving normal chow. After 14 days of treatment, mice were sacrificed, and tumors were harvested for enzymatic and mechanical digestion into single-cell suspensions. Whole tumor cell suspensions were profiled with scRNA-seq using 10x Genomics single cell Chromium library preparation. Both whole tumor and an immune-enriched single-cell suspension was created using magnetic bead selection for CD45+ cells and analyzed with single-cell ATAC-sequencing profiling using the sciATAC-seq technology.

The sciATAC-seq data were analyzed with Seurat (v 5.0.0)^64^ and Signac (v 1.12.9004)^65^ in R (v 4.3.1). Cells were retained if they had a nucleosome signal of less than 2 and a total number of peaks between 1000 and 8000 peaks. Cells with low transcription start site enrichment (< 1.5), high blacklist ration (>0.04) and low fraction of reads in peaks (< 0.3) were excluded from analysis. Seurat’s *FindNeighbors* and *FindClusters* functions were used for clustering and data visualization on UMAP space. Cells were initially annotated as cancer associated fibroblasts (CAF), Neoplastic, or Immune based on chromatin accessibility and Signac GeneActivity scores. To further define subpopulations of immune cells, scRNA-seq and sciATAC-seq were integrated using Seurat’s FindAnchors function for transfer learning^65,66^. Cell population labels were transferred from scRNA-seq to the sciATAC seq data using Canonical Correlation Analysis (CCA)^65^. Based on the labels from the scRNA-seq data, the annotation of immune cells in the sciATAC-seq data was able to be further refined into Lymphoid/NK and Myeloid cell types. The scRNA-seq data the sciATAC-seq data are available from NIH GEO.

TF activity scoring based solely on RNA expression data was conducted using pySCENIC^38^ (v 0.11.0). The list of genes encoding transcription factors and rankings of TF binding motif enrichment within 500 bp and 10 kb windows of gene transcription start sites in the mouse mm10 reference genome were obtained from the cisTarget^63^ resources website. Gene regulatory networks were learned from the raw RNA counts matrix using the grn function with method “GRNBoost.” Networks were pruned into TF regulons based on presence of TF binding motifs using the ctx function. TF activities were quantified in cells based on the learned regulons using the aucell function.

### Meta-cell integration of RNA expression and chromatin accessibility with SCENIC+

SCENIC+ requires clusters of cells and co-accessible genomic regions into regulatory topics as well as differentially accessible regions as input to identify candidate enhancers. The R package cisTopic (v 0.3.0)^67^ utilizes latent Dirichlet allocation to calculate the probability of a region belonging to a regulatory topic, and the probabilistic contribution of a regulatory topic within a cell. Counts from sciATAC-seq and cell metadata, including cell type annotation, were used to infer regulatory topics using cisTopic and converted to a pycisTopic object for compatibility with SCENIC+. pycisTopic (v 1.0.2) was used in python (v 3.11.9) to binarize the topics, impute accessibility, and infer differential accessibility between the cell types of interest using the binarize_topics, impute_accessibility, and find_diff_features functions, respectively SCENIC+ (v 1.0.1a) analysis was conducted using Snakemake (v 8.5.5) to quantify TF activity within cells based upon multi-omic RNA expression and chromatin expression profiles. As the scRNA-seq and sciATAC-seq data were collected in separate batches, pseudomulti-ome integration in SCENIC+ was used to integrate the scRNA-seq and sciATAC-seq based on cell type annotations in common across the modalities^39^. Shared annotations of cells as CAFs, Lymphoid/NK, Myeloid and Neoplastic cell types in the scRNA-seq and sciATAC-seq were used to create ‘meta-cells’ with SCENIC+. For each cell type, 10 cells of that type were randomly sampled from the scRNA-seq and sciATAC-seq data. A meta-cell was then created using the mean RNA expression and mean imputed accessibility of the sampled cells. The total number of meta-cells generated for each cell type was based on the minimum count of cells within each data modality and scaling this count by a scale factor defined in SCENIC+ to ensure an equal number of meta-cells were created for each data modality.

Using imputed chromatin accessibility from the meta-cells to specify putative enhancers, pycisTarget was used to identify differential motif enrichment between candidate enhancers and ranking-and-recovery of curated motifs maintained in the SCENIC+ collection database. GRNBooST2, as implemented in arboreto (v 0.1.6), was used to identify TF regulons based on co-expression of TFs and target genes and enrichment of TF motifs in enhancers of the target genes as identified by pycisTarget. TF regulon activity scores were quantified in cells using the AUCell function in the ctxcore (v 0.2.0) python package. Pseudomultiome meta-cells were cluster and visualized on UMAP based on TF regulon scores.

### DCST comparison between scRNA-seq analysis with SCENIC and integrated scRNA-seq and scATAC-seq data with SCENIC+

TF activity scores were calculated for CAF, Lymphoid/NK, Myeloid and Neoplastic cells in the UNT and ENT treatment groups were obtained from the RNA-seq data with SCENIC (v 0.11.0) and multi-omics data with SCENIC+ (v 1.0.1a) as described above. In order to assess the effects of using SCENIC or SCENIC+ on inference of intracellular linkages by dominoSignal, Fifteen bootstraps each were generated from profiles of cells in the UNT treatment groups of scRNAseq profiles (original RNA expression, SCENIC TF activity scores) and meta-cell profiles (meta-cell normalized RNA, SCENIC+ TF activity scores). Fifteen bootstraps were generated from UNT cells that underwent TF activity scoring with SCENIC or SCNEIC+. The number of bootstraps was chosen for computational efficiency. CCI was conducted using dominoSignal on each bootstrap as described in ‘Parameters for dominoSignal cell-cell communication inference on real data sets.’ For the TFs whose activity was inferred by both methods, intracellular linkages between these TFs and receptors were compared for all cell types by DCST using the bootstraps for SCENIC and SCENIC+. The number of intracellular linkages inferred between shared TFs for each cell type were plotted as a stacked barplot colored by whether the linkage was significantly differential in SCENIC, SCENIC+, or neither method. The overlap of inferred TF features for each method and treatment is shown as a Venn diagram generated using the ggvenn R package (v 0.1.10).

### Differential cell communication testing of effects of Entinostat on intercellular and intracellular communication in MMTV-PyMT tumors

Twenty bootstraps each were generated from the SCENIC+ meta-cells from the UNT and ENT treatment groups. Intercellular linkages for all recipient cell types were compared using DCST. Intercellular signaling involving macrophage inhibitory factor (*Mif*) were selected for further investigation as intercellular linkages that differentially occur between treatments based on *Mif* signaling having previously been demonstrated to impact survival outcomes in MMTV-PyMT tumors^43,44^. Tested Linkages involving the ligand *Mif* were compiled into one table and plotted using the dominoSignal differential signaling plot (Figure 6E). Intracellular linkages for each recipient cell types were compared using DCST. Linkages involving receptors of *Mif* (*Cxcr4*, *Cxcr2*, *Cd74*, *Cd44*, and *Ackr3*) were compiled, and an alluvial plot showing the significantly differential intracellular linkages between UNT and ENT was generated with R package ggalluvial (v 0.12.3) where the linkages are colored based on whether they were more likely to occur in UNT or ENT.

## Data and code availability

The dominoSignal package is available through Bioconductor (https://www.bioconductor.org/packages/release/bioc/html/dominoSignal.html). Scripts used for analysis presented in this manuscript are available from https://github.com/FertigLab/Differential_Cell_Signaling_Test.

## Acknowledgements

We would like to thank Dr. Michael Ochs and Dr. Atul Deshpande for providing feedback on the conceptualization of the DCST and bootstrapping methodologies. We would like to thank Wisam Awadallah, Tamara Lopez-Vidal, and Kylie Belanger for providing feedback on the user experience of dominoSignal. We would like to thank Dr. Peter Ordentlich of Syndax for providing the Entinostat utilized in this study. Figures 1, 3A, and 6B include graphics created using Biorender.

