## Supplementary figures and images for "Differential cell signaling testing for cell-cell communication inference from single-cell data by dominoSignal"

### Supplemental Figure S1

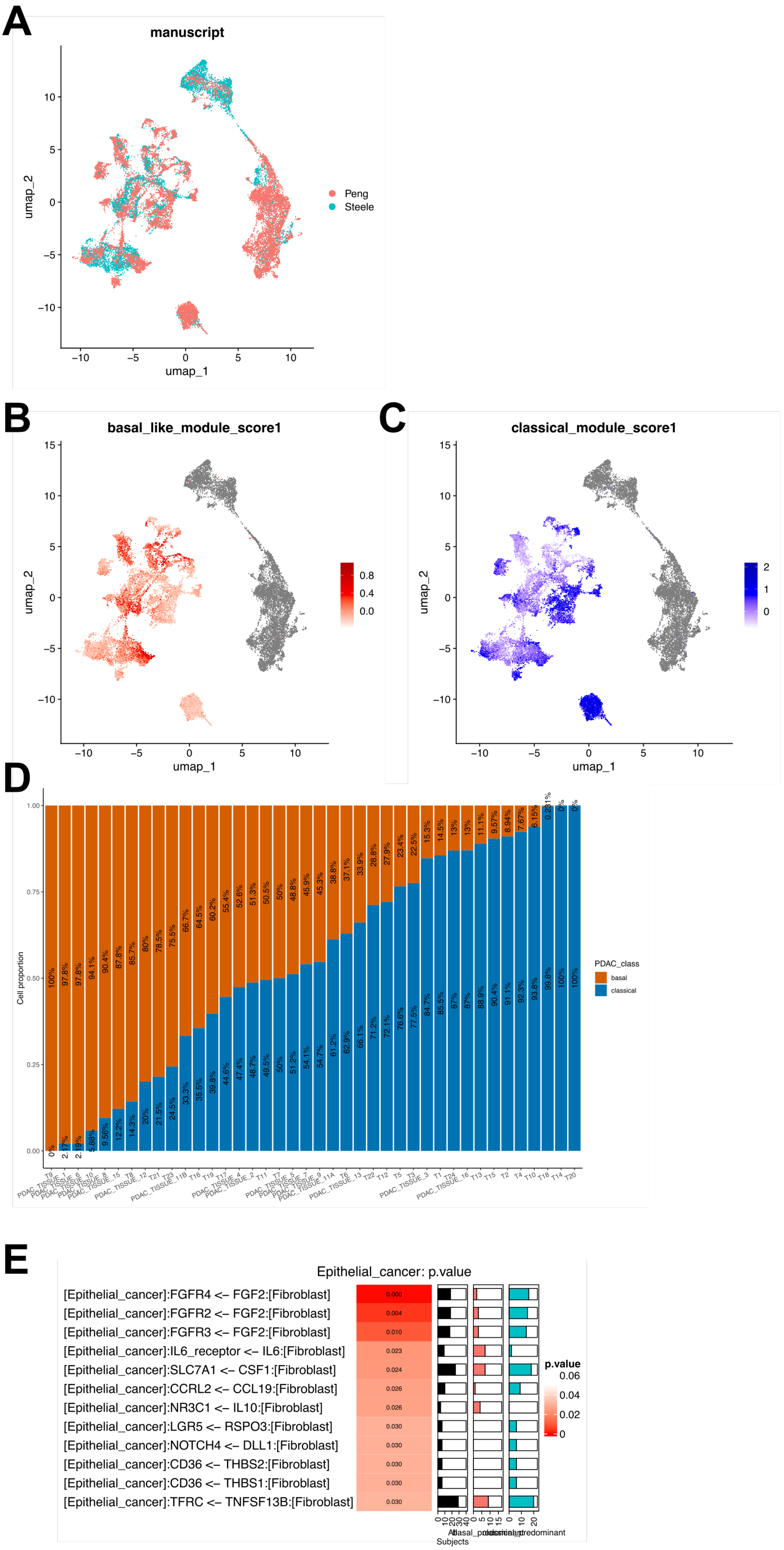

### Supplemental Figure S2

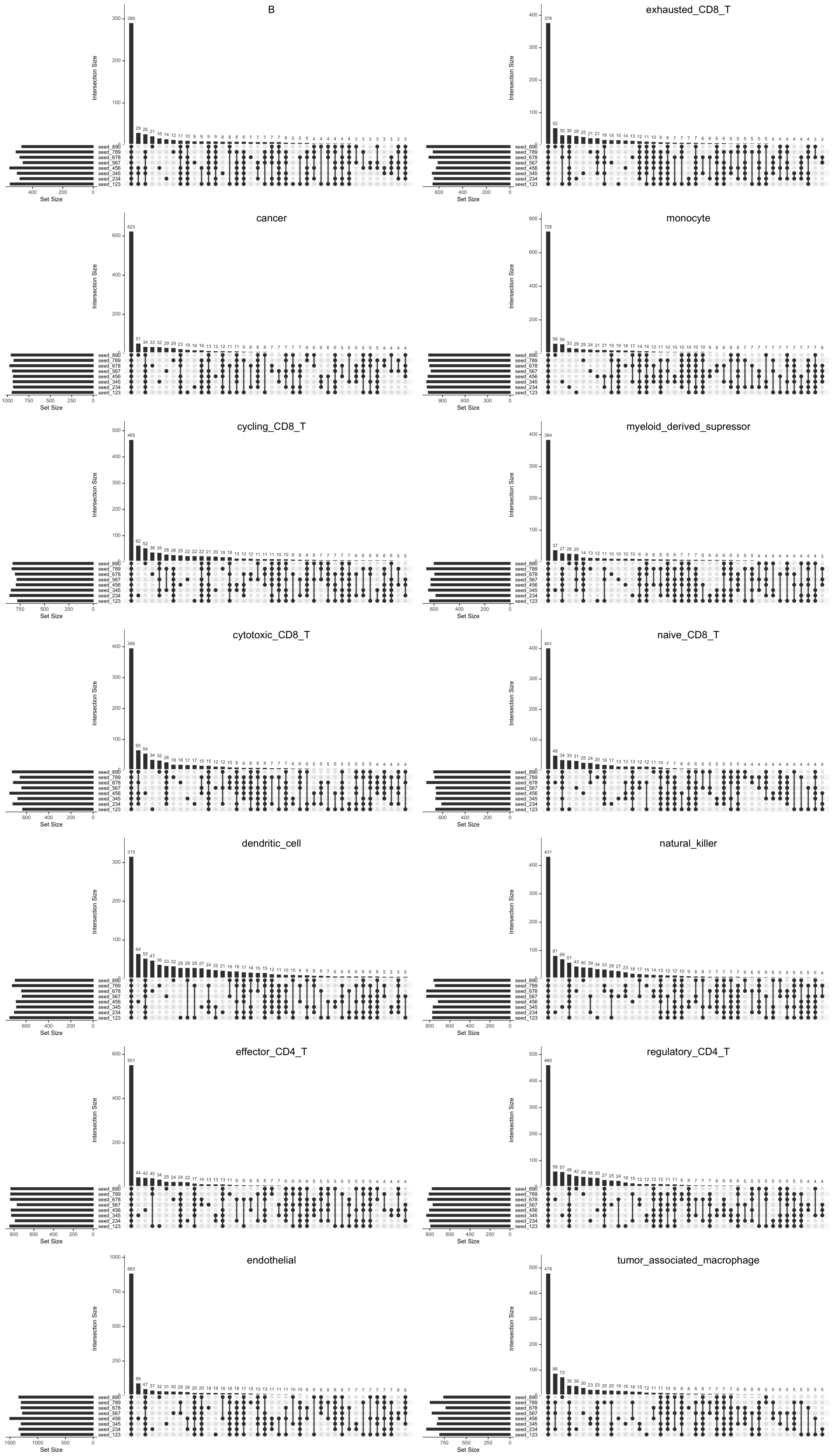

### Supplemental Figure S3

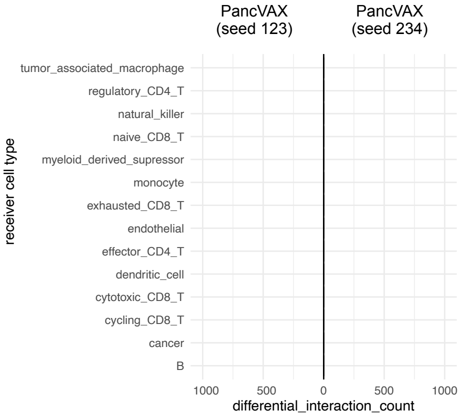

### Supplemental Figure S4

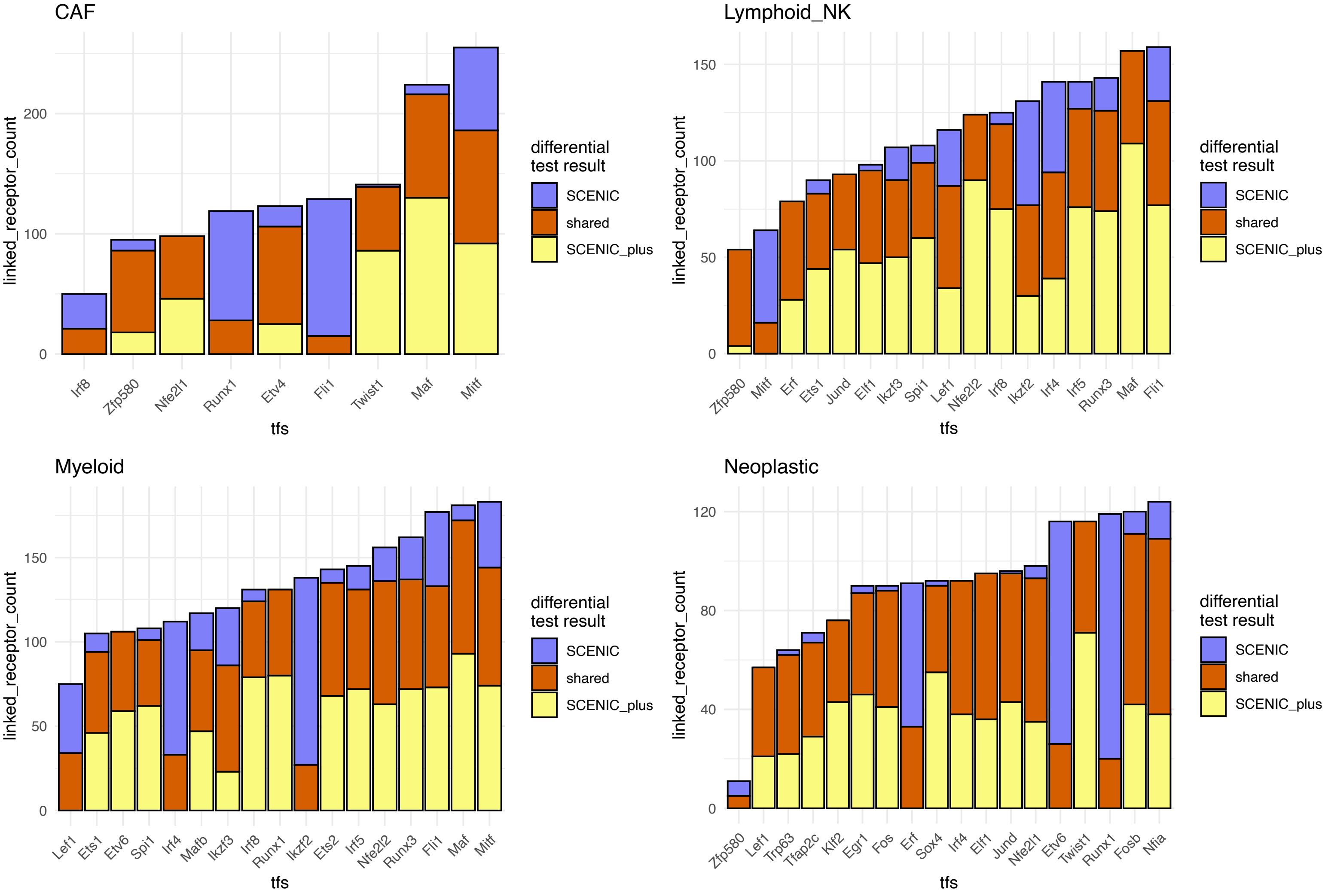
